# GAMETE maps the genetic architecture of chromatin accessibility in rice pollen at single-nucleus resolution

**DOI:** 10.64898/2026.07.04.736471

**Authors:** Yinmeng Liu, Chunjiao Xia, Junjie Li, Jiacheng Li, Haichuan Yang, Zhanxiang Zong, Luchang Ming, Junjiao Yang, Hantong Lian, Meng Luo, Tao Zhu, Xiaogang He, YongZhong Xing, Sibin Yu, Yidan Ouyang, Weibo Xie

**Author notes:** Correspondence: Weibo Xie.

## Abstract

**Background:** Dissecting the genetic basis of cell-type-specific gene regulation during gametogenesis remains a major challenge. Population-level studies mask cellular heterogeneity, while the extreme data sparsity of single-cell profiling has precluded robust genetic mapping.

**Results:** We present GAMETE (Genetic Architecture Mapping of Epigenetic Traits at single-nucleus resolution), an integrated framework combining MPS-ATAC-seq, a high-throughput single-nucleus ATAC-seq method, with robust computational approaches for genotype inference and genetic mapping from sparse single-nucleus data. Applying GAMETE to hybrid rice pollen, we simultaneously profiled chromatin accessibility and inferred the genotypes of 4,887 individual haploid nuclei, revealing distinct chromatin landscapes across pollen cell types. This uncovered an unexpected elevation of chromatin accessibility at *gypsy* retrotransposons in microspores preceding their silencing in sperm cells, exposing a surveillance gap that may contribute to genome expansion. The synchronized profiling enabled us to construct a single-cell recombination map, map segregation distortion loci and trace their developmental origins, and identify 16,113 chromatin accessibility QTLs (caQTLs), 48.6% of which exhibited pronounced cell-type specificity. Analysis of introgression lines supported a cell-type-specific *trans-*acting caQTL regulating the pollen-essential gene *DTM1*.

**Conclusions:** This work establishes GAMETE as a broadly applicable framework for dissecting the genetic architecture of chromatin accessibility during gametogenesis without the need for subsequent generations.

## Background

Cell-type-specific gene regulation during gametogenesis is essential for genome integrity and reproductive success. Yet how genetic variation shapes these regulatory programs remains poorly understood. Pollen development in F₁ hybrids offers a powerful system to dissect this complexity, as it simultaneously generates functionally distinct haploid lineages and naturally segregates parental genomes. In higher plants, meiosis of pollen mother cells produces haploid microspores (MC), which develop through asymmetric division into pollen grains containing a vegetative cell (VC) and a generative cell (GC)^1^. The GC further divides to form two sperm cells (SC)^2^. Although this precisely regulated developmental program is essential for plant fertility, the genetic basis of cell-type-specific chromatin regulation during pollen development remains largely unexplored. For example, while VC are known to produce small RNAs that non-cell-autonomously silence transposons in SC via RNA-directed DNA methylation (RdDM)^3,4^, population genomics reveals pervasive transposon insertion polymorphisms (TIPs) across rice accessions^5^, suggesting that transposon regulation during pollen development may be more dynamic than implied by the canonical silencing model.

Resolving cell-type-specific genetic regulation in pollen faces three major technical challenges. First, genetic mapping of pollen-related traits typically requires evaluation in subsequent generations, severely limiting research efficiency. For instance, identifying segregation distortion loci in F_1_ pollen traditionally requires growing and genotyping hundreds of F_2_ plants^6^. Second, molecular studies demand comprehensive omics data, which is challenging and costly to acquire for pollen due to its limited availability and transient developmental windows. Third, the asynchronous nature of pollen development introduces sample heterogeneity, while the presence of distinct cell types within individual pollen grains masks cell-type-specific chromatin states even in single-pollen analyses^7^. Although single-cell profiling can in principle resolve this heterogeneity, the extreme data sparsity of current approaches has precluded reliable genetic mapping from individual cells.

To address these challenges, we developed GAMETE (Genetic Architecture Mapping of Epigenetic Traits at single-nucleus resolution), an integrated framework comprising three components: (1) Multiplex Plate-based Single-nucleus ATAC-seq (MPS-ATAC-seq), a high-throughput platform for single-cell chromatin accessibility (CA) analysis; (2) a Hidden Markov Model (HMM)-based approach^8^ for accurate genotype inference from sparse single-cell data; and (3) a Dynamic Bulk Segregant Analysis (Dynamic-BSA) method that leverages recombinant genotypes in gametes for efficient quantitative trait locus (QTL) mapping. Applying GAMETE to hybrid rice Shanyou 63 (SY63) pollen yielded unexpected insights into epigenetic regulation and genetic architecture. Analysis of 4,887 high-quality haploid nuclei uncovered distinct chromatin landscapes across cell types and, surprisingly, elevated CA at retrotransposons in MC. By exploiting natural genetic variation in F_1_ hybrid pollen, we traced the developmental origins of segregation distortion and identified 16,113 CA QTLs (caQTLs), nearly half with pronounced cell-type specificity. Integration with deep learning further enabled the prediction of regulatory effects of genetic variants. Beyond pollen, GAMETE establishes a broadly applicable framework for dissecting the genetic architecture of CA during gametogenesis across species.

## Results

### MPS-ATAC-seq enables flexible single-nucleus chromatin accessibility profiling

Single-cell ATAC-seq (scATAC-seq) has become a pivotal technique for identifying cell-type-specific regulatory elements across diverse organisms^9,10,11^. Among the various scATAC-seq approaches, the microfluidic-based system from 10X Genomics Chromium has gained widespread acceptance^12,13^. However, its application is largely limited to commercial use, constrained by high costs and inconvenience. More recently, a cost-effective and flexible plate-based scATAC-seq has been developed^14^. Building upon this foundation, we have developed a more flexible indexing strategy, resulting in a method we term multiplex plate-based single-nucleus ATAC-seq (MPS-ATAC-seq).

The workflow of MPS-ATAC-seq is summarized as follows (Fig. 1a): First, pollen nuclei were isolated from SY63 during the flowering and pollination stages. Fluorescence-activated cell sorting (FACS) was employed to perform the first round of sorting, resulting in the collection of 24 tubes, each containing approximately 30,000 nuclei. Next, to distinguish the nuclei in the 24 tubes, unique barcodes were added to the adapters of the Tn5 transposase, with each tube receiving a distinct barcode. This was followed by 24 independent transposition reactions, ensuring that the nuclei in each tube were labeled with unique adapters. Subsequently, the 24 tubes of nuclei were split into 384-well plates. As a result, each well contained 24 nuclei, each labeled with a distinct Tn5 barcode (Methods). In total, this approach theoretically allows the profiling of up to 9,216 nuclei (24 × 384) in a single MPS-ATAC-seq experiment (Fig. 1a, Methods).

**Fig. 1:**
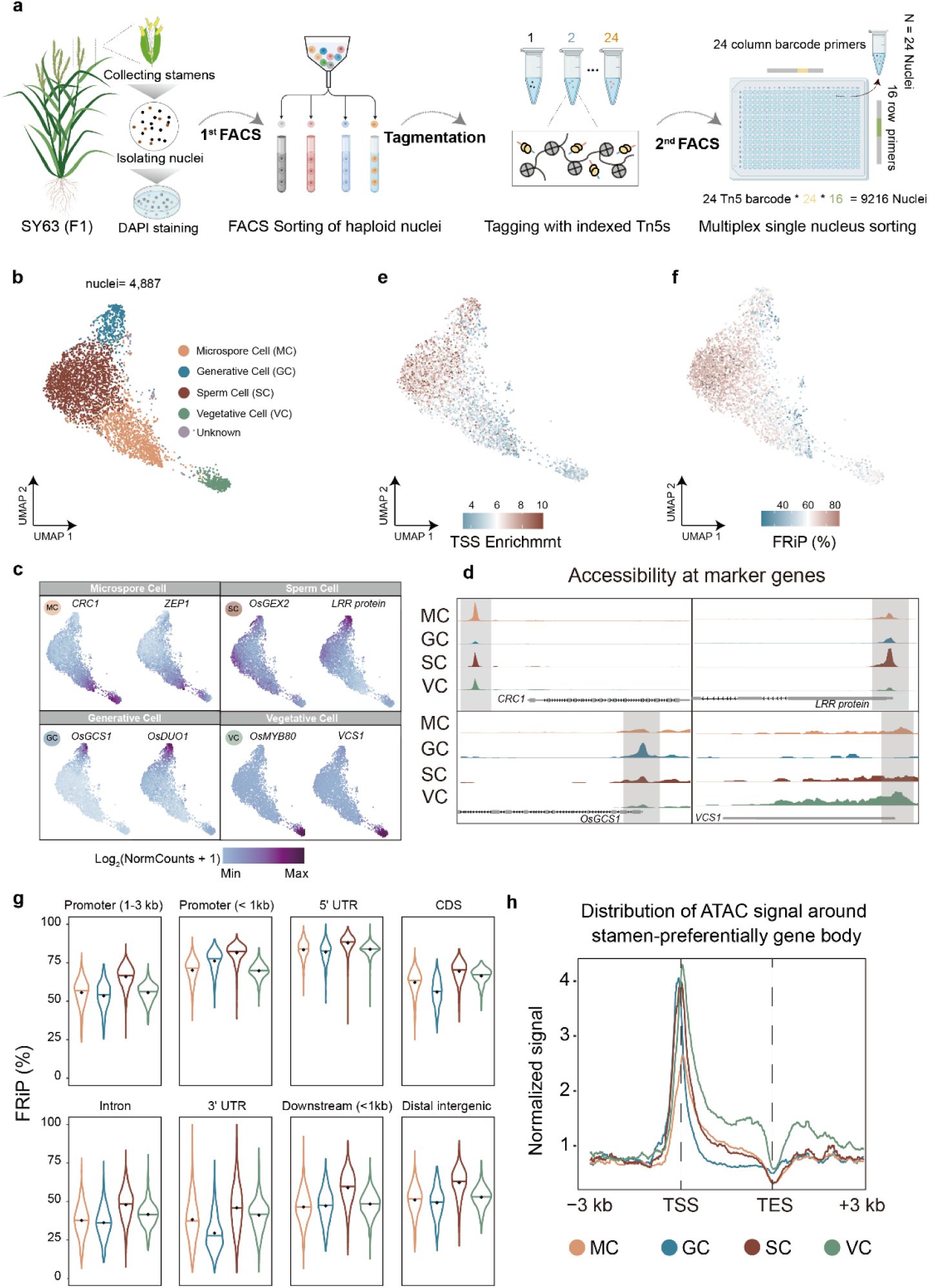
Overview of the MPS-ATAC-seq experimental workflow and data quality. **a**, Schematic of the MPS-ATAC-seq experimental workflow. Pollen nuclei were isolated and sorted by fluorescence-activated cell sorting (FACS), followed by indexed Tn5 tagmentation and multiplex single-nucleus sorting into 384-well plates. **b**, UMAP projection of nuclei colored by major clusters. Each point represents a single nucleus, with different colors denoting different cell types. Four main cell types were identified: MC (microspore cell), GC (generative cell), SC (sperm cell), and VC (vegetative cell). **c**, Chromatin accessibility (CA) patterns at regulatory regions of cell-type-specific marker genes. **d**, Aggregated CA profiles surrounding known marker genes across different cell types. **e-f**, UMAP projection of nuclei colored by transcription start site (TSS) enrichment scores and the fraction of reads in peaks (FRiP) values among different cell types. Cell type annotations correspond to panel **b**. **g**, Distribution of FRiP scores across different genomic annotations for each cell type. Genomic regions are categorized as follows based on their positions: promoter 1-3 kb (1 to 3 kb upstream of promoters), promoter <1 kb (within 1 kb upstream of promoters), 5′ UTR (5′ untranslated region), CDS, intron, 3′ UTR (3′ untranslated region), downstream < 1 kb (within 1 kb downstream of genes), and distal intergenic (encompassing all remaining genomic regions not classified above). The y-axis represents the percentage of reads mapped to accessible chromatin regions (ACRs) overlapping with each specific genomic feature. In the violin plots, the horizontal line represents the median values, and the black dot represents the mean values. **h,** Distribution of whole-genome normalized CA within 3 kb upstream and downstream of stamen-preferentially genes (2,586 genes) across different cell types.

### MPS-ATAC-seq unveils cell-type-specific chromatin landscapes during rice pollen development

We performed two biological replicates using the MPS-ATAC-seq method. Following sequencing, nuclei were filtered based on stringent quality control criteria. In addition to standard metrics such as Tn5 insertions and transcription start site (TSS) enrichment scores, we applied specific filters for genomic coverage uniformity (chi-square test) and genotype consistency (correlation coefficient) to exclude low-quality or contaminated nuclei (Additional file 1: Fig. S1a-e, Methods). After filtering, 4,887 high-quality nuclei were retained for downstream analysis, with an average of 2,616 unique Tn5 insertions per nucleus (Fig. 1b, Additional file 1: Fig. S1c, d, Methods). The aggregated CA profiles from these nuclei exhibited high concordance with bulk ATAC-seq data, confirming the reliability of our single-nucleus dataset (Additional file 1: Fig. S1e).

Using the ArchR package^15^, we performed dimensionality reduction and unsupervised clustering, revealing four major clusters. To assess data quality and reproducibility, we examined potential batch effects between replicates and found uniform distribution of nuclei across clusters, confirming minimal technical variation (Additional file 1: Fig. S1g, h). We therefore merged the datasets for subsequent analyses. Based on CA patterns of established marker genes (Fig. 1c,d, Additional file 1: Fig. S2b, Methods), we annotated these clusters as nuclei from microspore cells (MC, n=1,466), generative cells (GC, n=524), sperm cells (SC, n=2,330), and vegetative cells (VC, n=478), with a small fraction from unidentified cell types (n=89) (Fig. 1b, Methods). Expression patterns derived from published rice pollen RNA-seq datasets^16,17^ were broadly concordant with the CA profiles of representative cell-type marker genes (Additional file 1: Fig. S2g). The predominance of SC nuclei and the smaller proportions of VC and GC nuclei in our dataset reflect the cellular composition of pollen at the collection stage. We also note that VC may be more susceptible to damage during isolation, resulting in their underrepresentation (Additional file 1: Fig. S1g, Methods).

To further characterize the CA landscape of each cell type, we identified 53,619 accessible chromatin regions (ACRs) covering 7.8% of the rice genome from an independent SY63 pollen bulk ATAC-seq dataset, which served as an unbiased reference for quality assessment (Methods). The four cell types differed markedly in overall CA levels, with VC showing the highest and SC the lowest number of uniquely mapped Tn5 insertions (Additional file 1: Fig. S1j). Despite having the fewest insertions, SC exhibited the highest TSS enrichment scores and FRiP values among all cell types (Fig. 1e,f, Additional file 1: Fig. S1k, l). To resolve these differences at finer resolution, we categorized the genome into eight regions and compared FRiP scores and read distributions within each across the four cell types (Fig. 1g, Additional file 1: Fig. S1m). SC exhibited the highest FRiP values across nearly all regions (Fig. 1g). In “Promoter < 1kb” regions, VC and MC showed notably lower FRiP scores than GC and SC. Across genic regions (5 ′UTR, CDS, intron, and 3′ UTR), GC consistently displayed the lowest FRiP scores, indicating a broad reduction of accessibility across gene bodies. In contrast, VC showed relatively uniform FRiP values across all genomic regions (Fig. 1g). Together, these results indicate that CA in SC is highly concentrated at specific regulatory elements, whereas CA in VC, despite being the most abundant overall, is distributed broadly across the genome. Consistently, VC displayed higher CA than SC and GC at gene body and downstream regions of stamen-preferentially expressed genes (Fig. 1h). These patterns were reproducible across biological replicates (Additional file 1: Fig. S1i).

Using a max gap-based approach, we identified 5,468 ACRs exhibiting pronounced cell-type-preferential accessibility (Additional file 1: Fig. S2a, b, Additional file 2: Table S5, Methods). Pattern analysis revealed that GC undergo the most extensive chromatin remodeling, with over half of these ACRs showing either GC-preferential activation (30.65%) or repression (24.56%). The second major pattern was VC-preferential activation (20.30%). GO analysis revealed that genes associated with GC-activated ACRs were enriched in transcription regulation and nucleic acid metabolic process, while genes associated with VC-activated ACRs were enriched in metabolic enzymes including amylase activity, reflecting regulatory versus metabolic specialization (Additional file 1: Fig. S2c, d). Representative genes from these categories showed cell-type-preferential expression in published pollen transcriptomes, consistent with their CA (Additional file 1: Fig. S2f, g).

### Microspores exhibit elevated chromatin accessibility at retrotransposons

Previous research has demonstrated differential transposable element (TE) activity between cell types, with silenced TEs becoming reactivated in VC while remaining repressed in SC^3^. To investigate TE accessibility patterns across pollen cell types, we collected rice TE datasets^18^ and quantified CA at TEs by calculating the proportion of sequencing reads mapping to TEs relative to total mapped reads (Fig. 2a, Methods).

**Fig. 2:**
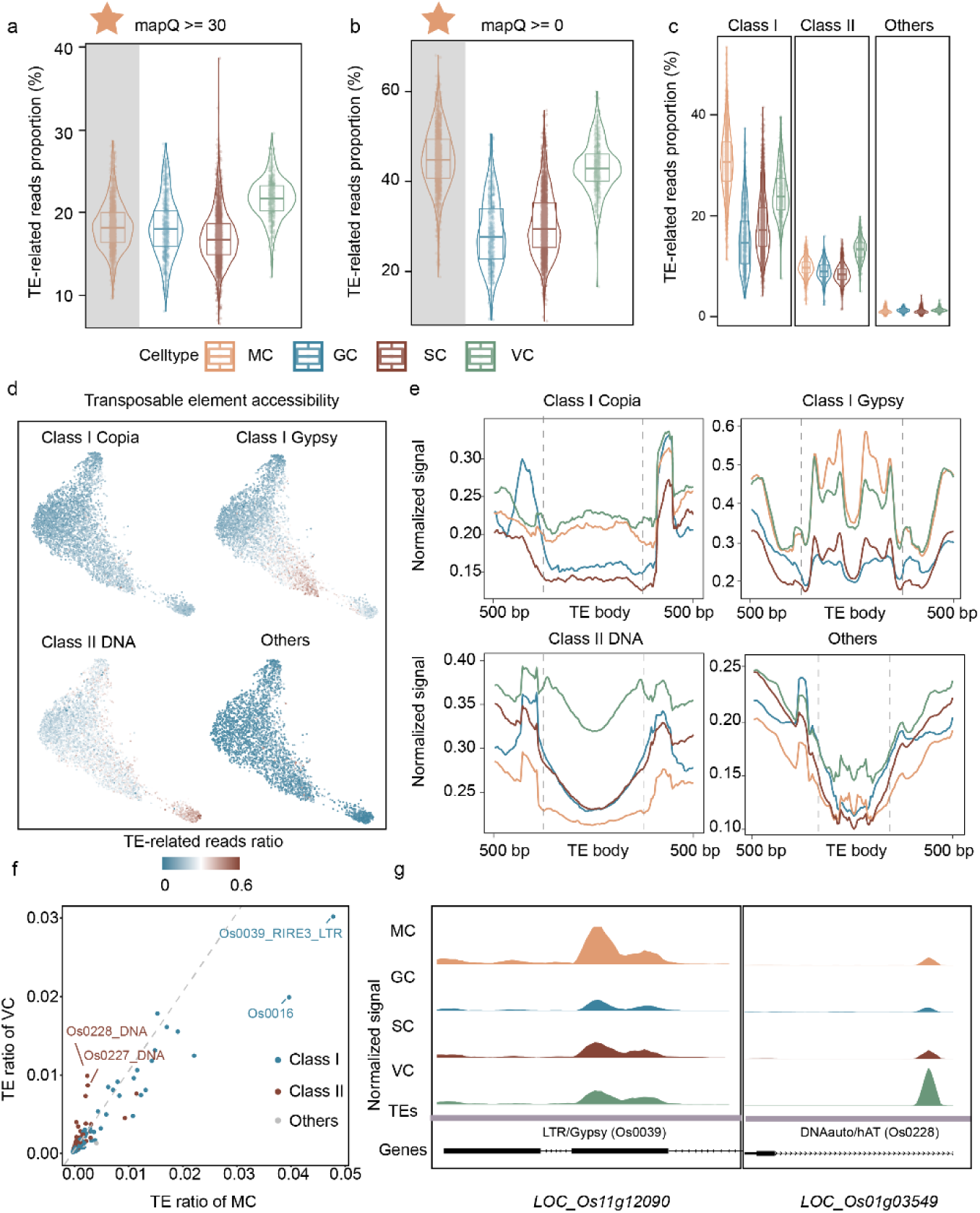
Landscape of transposable element (TE) accessibility across cell types. **a-b**, Percentage of reads mapping to TEs across different cell types. Reads were filtered based on MAPQ scores ≥ 30 **(a)** and ≥ 0 **(b)** using SAMtools, respectively. The y-axis represents the proportion of TE-mapped reads relative to the total reads in each nucleus. **c**, Chromatin accessibility (CA) of three major TEs categories [Class I (retrotransposons), Class II (DNA transposons) and others] across different cell types. **d**, UMAP projection showing CA of TEs across different cell types. Color intensity indicates the proportion of reads mapping to TEs (darker colors indicate higher proportions). **e**, Distribution of CA around TEs across different cell types. The lengths of Class I and Class II transposons were normalized to 1 kb, while the rest were normalized to 500 bp. **f**, Comparison of the proportions of reads mapping to different TE subfamilies between MC and VC. Each dot represents a TE subfamily. The dashed line indicates equal proportions. Representative subfamilies deviating from this diagonal are labeled. **g**, Genome browser tracks showing CA at representative TE loci from two different subfamilies, LTR/Gypsy (Os0039, left) and DNAauto/hAT (Os0228, right), across cell types.

Our analysis using standard data processing pipelines revealed higher CA at TEs in VC compared to SC, consistent with previous findings in *Arabidopsis*^3^. However, we recognized that standard pipelines (e.g., using SAMtools with a MAPQ ≥ 30 threshold)^19^ often exclude alignments derived from repetitive sequences. Given that TEs typically exist as multiple genomic copies, applying such filtering may inappropriately exclude TE-derived reads. Therefore, we analyzed data without filtering alignment quality scores. The results showed not only the expected VC-SC difference but also unexpectedly high TE accessibility in MC (Fig. 2a-b), a phenomenon not previously reported.

To gain deeper insights into TE-associated chromatin dynamics during pollen development, we systematically categorized TEs into Class I (mainly retrotransposons), Class II (mainly DNA transposons), and others, and analyzed their accessibility across cell types. The results revealed that Class II transposons exhibited higher CA in VC compared to other cell types (Fig. 2b), primarily contributed by the DNA transposon subtypes DNA/MULE, DNA/CACTG, and DNA/hAT (Fig. 2c-d). In contrast, Class I TEs showed significantly higher CA in MC compared to GC, SC, and VC (Fig. 2c), with the major contributors being *gypsy*-type retrotransposons (Fig. 2d, Additional file 1: Fig. S3a). To further characterize the chromatin landscape, we examined accessibility signals at and flanking TE bodies by scaling TE body lengths to a uniform length (Fig. 2e). Both MC and VC exhibited elevated CA at *gypsy* retrotransposon bodies and flanking regions relative to GC and SC, with MC showing the highest levels. In contrast, elevated CA at DNA transposons was specific to VC (Fig. 2e). Comparative analysis of the proportions of reads mapping to different TE subfamilies between MC and VC further supported this cell-type-specific partitioning (Fig. 2f). Within MC, the elevated TE accessibility was predominantly driven by two *gypsy* subfamilies, Os0039_RIRE3 and Os0016 (Additional file 1: Fig. S3a-c). Strikingly, these same two subfamilies were recently identified as the primary contributors to genome size divergence between wild and cultivated rice, having undergone the most pronounced copy number contraction during domestication^20^ (Additional file 1: Fig. S3d). In contrast, VC showed increased accessibility at specific DNA transposon subtypes, particularly hAT and MULE elements (Fig. 2f,g).

Together, these results reveal a previously undescribed cell-type-specific partitioning of TE accessibility during pollen development. Retrotransposon accessibility is elevated in both MC and VC relative to the germline cells, but is most pronounced in MC, whereas elevated DNA transposon accessibility is specific to VC. The convergence between the *gypsy* subfamilies showing the highest accessibility in MC and those driving the most pronounced genome size changes during domestication^20^ suggests that the transient accessibility window in MC may provide an opportunity for retrotransposon amplification, potentially influencing genome size evolution.

### Dynamic chromatin accessibility underlies developmental trajectories

To investigate the developmental progression of rice pollen, we performed pseudotime analysis on MC, GC, and VC, excluding SC as it differentiates from GC rather than directly from MC. Trajectory inference based on 4,268 cell-type-specific ACRs across 2,468 nuclei revealed a bifurcating developmental path from MC, with one branch leading to GC and the other to VC (Fig. 3a-c).

**Fig. 3:**
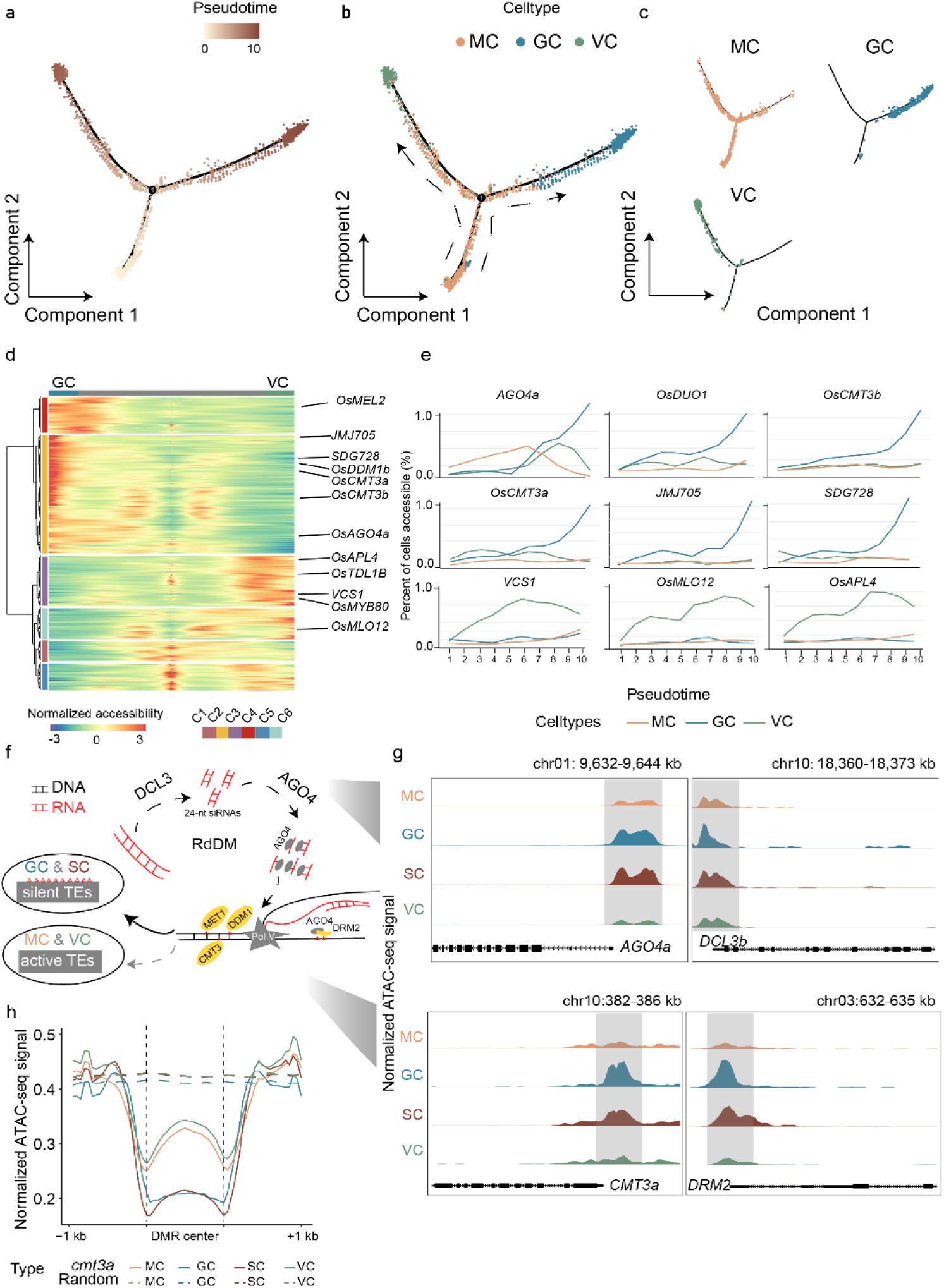
Dynamic chromatin accessibility changes during rice pollen development. **a-c,** UMAP projections showing the developmental trajectory of rice pollen. Cells are colored by pseudotime (**a)** and cell types (**b-c)**. **d**, Heatmap of chromatin accessibility (CA) depicting six clusters of significantly differential accessible chromatin regions along the MC-GC and MC-VC pseudotime trajectories. **e**, Line plots illustrating CA dynamics of nine representative genes along MC-GC and MC-VC differentiation branches. Line colors correspond to respective cell types. **f**, Schematic model of the RdDM pathway and the epigenetic regulation of transposable elements across different rice pollen cell types. **g**, Normalized ATAC-seq signals at regulatory regions of key RdDM pathway genes across MC, GC, SC, and VC cell types. **h**, Normalized ATAC-seq signals at CHG-hypomethylated DMRs (identified in *cmt3a* mutant sperm cells) across different cell types. Randomly selected genomic regions serve as background controls

We identified 2,000 genes associated with ACRs exhibiting significant accessibility changes along the pseudotime trajectory (Fig. 3d), demarcating divergent developmental programs. Along the MC-to-GC branch, *OsDUO1* (ortholog of *Arabidopsis DUO1*, a master regulator of male germline specification) showed progressively increased accessibility^21^. The H3K27me3 demethylase *JMJ705* displayed progressive accessibility increase along the GC trajectory, aligning with findings that H3K27me3 removal is required for male germline fate initiation while its maintenance is essential for VC commitment^22^. Along the MC-to-VC branch, *VCS1* (vegetative cell-specific marker)^23^, *OsMLO12* (seven-transmembrane protein essential for pollen hydration and germination)^24^, and *OsAPL4* (large subunit of ADP-glucose pyrophosphorylase required for starch synthesis)^25^ displayed increased accessibility, reflecting the specialized metabolic and developmental functions of VC (Fig. 3e).

Given the prominent differences in TE-associated CA between cell types, we examined CA of the RNA-directed DNA methylation (RdDM) pathway components (Fig. 3f). Analysis revealed striking cell-type-specific patterns (Fig. 3g). Multiple genes involved in TE silencing, including *DCL3b* (stamen-specific 24-nt phased small RNAs)^26^, *DRM2* (de novo methyltransferase)^27^, and *AGO4a* (the most accessible *AGO* gene in pollen)^28,29^, exhibited elevated accessibility in GC and SC but reduced accessibility in MC and VC. Since *AGO4a* binds 24-nucleotide siRNAs produced by *DCL3* and guides *DRM2*-mediated DNA methylation to TEs, the coordinated increase in accessibility of these RdDM pathway components in GC and SC is consistent with enhanced genome protection in the male germline, while their reduced accessibility in MC and VC might contribute to observed TE-associated CA dynamics in these cell types.

Furthermore, chromatin remodeling, histone modification, and DNA methylation maintenance enzymes displayed coordinated lineage-specific dynamics reinforcing these patterns. *OsDDM1b*, a chromatin remodeling factor essential for maintaining DNA methylation and stable heterochromatin^30^, exhibited markedly increased accessibility in GC but decreased accessibility in VC. *SDG728*, an H3K9 methyltransferase^31^, and *CMT3a*/*CMT3b*, which recognize H3K9me2 to catalyze CHG methylation, showed similar patterns with increased accessibility in GC but reduced accessibility in VC^16^. *OsMET1b*, the maintenance CG methyltransferase^32^, also exhibited increased accessibility in GC but reduced accessibility in VC (Fig. 3d), suggesting potential differences in CG methylation maintenance capacity between lineages. Among the RdDM and related silencing components examined above, those expressed in the germline showed expression patterns consistent with their accessibility (Additional file 1: Fig. S2g). These patterns suggest that both de novo and maintenance methylation pathways are coordinately accessible during the MC-to-GC transition, while remaining low in the MC-to-VC trajectory, which may contribute to the observed differences in TE chromatin state among cell types.

We next examined whether DNA methylation is associated with these CA patterns. Since CHG methylation is a major silencing mark at plant retrotransposons and is primarily deposited by CMT3-type chromomethylases, we examined published bisulfite sequencing data from *cmt3a* mutant sperm cells ^16^. Regions that lose CHG methylation in SC of *cmt3a* mutants (indicating *CMT3a* targets) exhibited lower accessibility in GC and SC compared to MC and VC (Fig. 3h). This links CMT3-mediated CHG methylation to cell-type-specific CA dynamics and suggests that these CHG-silencing targets are less suppressed in MC than in the male germline.

Together, these results suggest that rice pollen development involves coordinated, cell-type-specific epigenetic reprogramming: the distinctively low accessibility of RdDM and related silencing components in MC correlates with the observed elevated CA at retrotransposons, while their increased accessibility in GC and SC may contribute to genome stability in the male germline, paralleling the establishment of specialized chromatin states in VC.

### Construction of the genotype recombination map

SY63 is an F_1_ hybrid derived from a cross between two inbred parental lines MH63 and ZS97. As a result of homologous recombination during meiosis in pollen mother cells, each resulting pollen grain harbors a unique mosaic genotype with chromosomal segments inherited from both parental genomes. By analyzing the sequencing data from individual pollen nuclei, it is possible to obtain genotype information, enabling the construction of a high-resolution genetic recombination map. However, each nucleus has an average of only 20.24% reads covering polymorphic sites between the two parental varieties. This sparse coverage, combined with potential exogenous DNA contamination during single-nucleus isolation and inevitable sequencing errors, poses significant challenges to constructing a reliable genotype recombination map.

To mitigate these potential errors, we adapted an HMM based on our previous research^8^, effectively inferring genotypes for individual nucleus by integrating information such as the physical distance of variant sites, recombination rates, and sequencing error rates (Fig. 4a-b, Methods). We detected 102,786 recombination events in 4,887 nuclei, with an average of approximately 21 recombination events per gamete (Additional file 1: Fig. S4a). Recombination was less frequent near centromeric regions, consistent with the recombination distribution previously identified from MH63 and ZS97 recombinant inbred line populations (Additional file 1: Fig. S4b)^33^. High-recombination genomic regions exhibited significantly higher CA than the rest (Wilcoxon rank-sum test, *P* =2.9 × 10^−4^; Additional file 1: Fig. S4c).

**Fig. 4:**
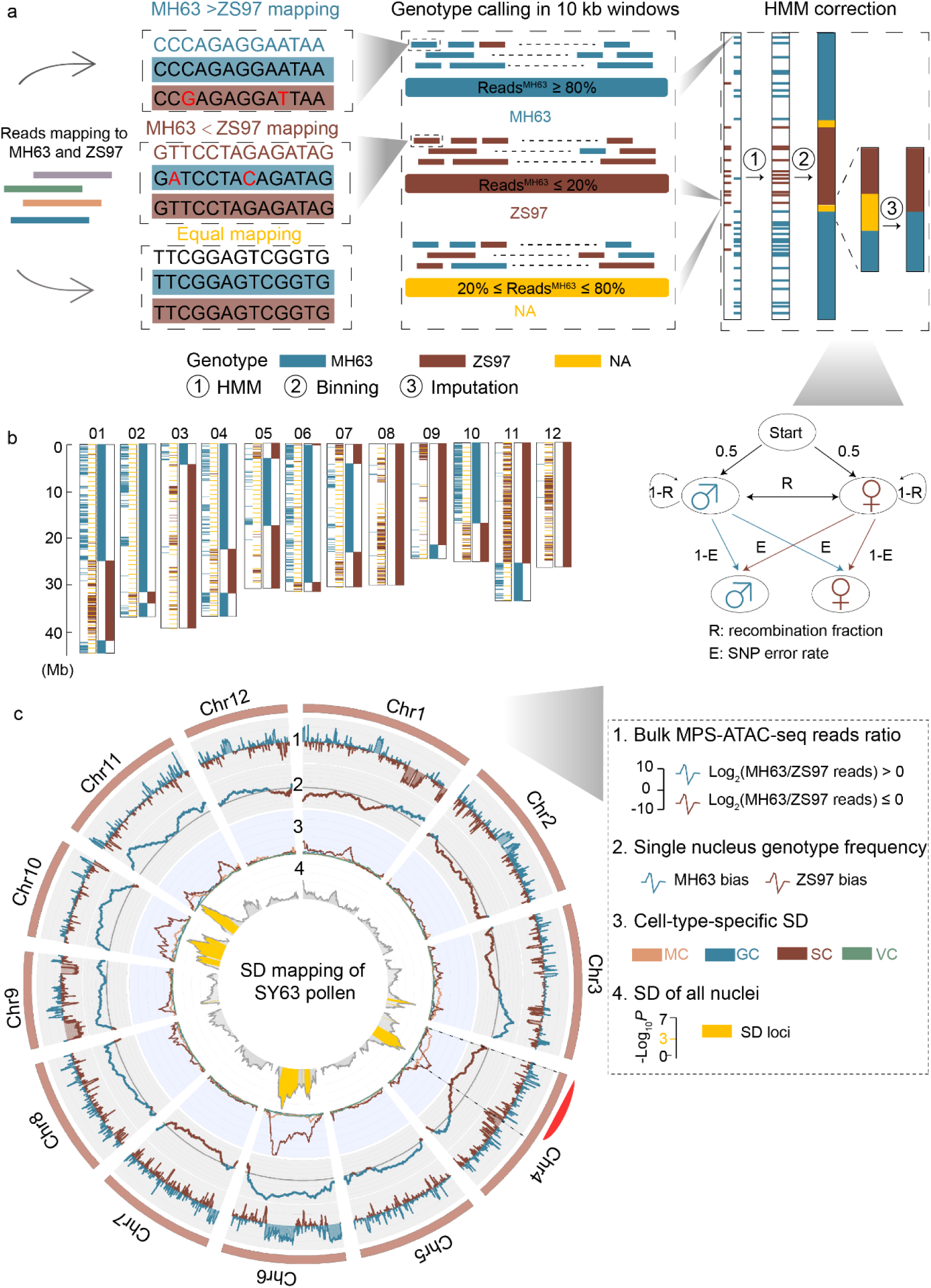
Genotype map construction and segregation distortion mapping. **a**, Workflow for constructing the genotype map. **b**, Example of genotype determination in a randomly selected nucleus, illustrating the processing steps: raw genotype calling, HMM correction, and final imputation. Blue regions indicate MH63 genotypes and brown regions indicate ZS97 genotypes. **c**, Identification of segregation distortion (SD) loci in SY63 pollen, shown from outer to inner panels. To ensure data uniformity for visualization, the plotted values represent the mean of 10 kb analysis windows averaged over 100 kb intervals: **1**, Bulk reads ratio from MPS-ATAC-seq data. The y-axis represents the log_2_ ratio of MH63 to ZS97 reads. Positive values indicate a bias towards MH63 (depicted in blue), while negative values indicate a bias towards ZS97 (depicted in brown). **2**, Genotype frequency calculated by MPS-ATAC-seq data. For each 10 kb window, nuclei were classified as MH63 or ZS97, and the proportion of each genotype across all nuclei was calculated. Windows with MH63 proportion >0.5 are considered biased toward MH63 (depicted in blue), while segments with a ratio <0.5 are considered biased toward ZS97 (depicted in brown). **3**, Cell-type-specific SD loci identified using MPS-ATAC-seq data. For each cell type, a chi-square test was performed within each 10 kb window to test for deviation from the expected 1:1 genotype ratio. The y-axis shows the −log_10_(*P* value). **4**, SD across all nuclei using MPS-ATAC-seq data. For each 10 kb window, genotypes were classified as MH63 or ZS97, and a chi-square test was conducted across all nuclei to genotype counts against the expected 1:1 ratio. The y-axis represents the −log_10_ (*P* value) of the chi-square test. Regions with −log_10_ (*P* value) >3 are defined as SD loci and highlighted in yellow. The red arcuate region indicates the SD loci identified in a previous study^6^.

Despite the low sequencing coverage typical of single-nucleus datasets, our method yielded reliable genotypes, providing a robust foundation for subsequent analyses. Unlike traditional methods requiring F_2_ populations and conducting extensive offspring resequencing, our study provides a more direct and efficient alternative approach for mapping the recombination landscape directly from F_1_ pollen.

### Map segregation distortion loci and trace their developmental origins

Segregation distortion (SD), the deviation of allele frequencies from expected Mendelian ratios, is a widespread phenomenon with significant implications for plant breeding and evolution. Using the constructed genotype map, we performed chi-square tests to detect SD and identified eight loci with significant deviations from expected segregation ratios^34^ (Fig. 4c, Methods). Most loci showed a bias toward MH63, except for one on chromosome 4, which displayed a bias toward ZS97 with a segregation ratio of 0.54 (Fig. 4c, Methods). Previous studies using an F_2_ population (n=163) from a ZS97 × MH63 cross identified two SD loci, including one on chromosome 4 that overlaps with our findings. Consistent with previous findings, the genotype at this locus also showed a bias toward ZS97, confirming the accuracy of our SD identification. While the earlier study localized this locus to a 0-19.82 Mb interval^6^, our approach has narrowed it down to 12.68-20.99 Mb, substantially enhancing the precision of the SD detection and providing valuable insights for future gene mapping. Notably, a large inversion (18.96-20.06 Mb) between MH63 and ZS97 has been reported in this region of chromosome 4^35^ (Additional file 1: Fig. S5). This structural variation may underly the observed SD, though the exact mechanism requires further investigation.

By integrating our nucleus type annotations with the genotype data, we performed stage-specific segregation analysis. For each annotated nucleus population (MC, GC, SC, and VC), we separately conducted chi-square tests to compare observed genotype frequencies against expected Mendelian ratios. This approach enabled us to track SD loci across the developmental continuum of pollen maturation. Notably, SD loci in the central portion of chromosome 4 and the proximal end of chromosome 11 were detected as early as the MC stage (Fig. 4c), providing direct evidence for the timing of these genetic aberrations. In summary, our approach of leveraging single-cell sequencing data enabled us to precisely identify SD loci and to trace the developmental origins of SD loci, overcoming the limitations of traditional genetic population-based analyses that cannot directly access pre-fertilization stages or determine when transmission barriers first emerge, thus opening new avenues for understanding genetic transmission barriers in plants.

### Dynamic-BSA enables identification of cell-type-specific chromatin accessibility QTLs

CA at individual ACRs might vary among nuclei and can be viewed as a quantitative trait associated with genotypes. While QTL mapping approaches should theoretically enable dissection of the genetic basis of CA, the sparsity of single-nucleus data poses significant challenges for traditional methods.

To address this limitation, we developed Dynamic-BSA. Unlike traditional BSA, which forms bulks by phenotype^36^, it dynamically reassigns nuclei into genotype-defined bulks at each genomic interval, enabling CA to be mapped as a quantitative trait. Briefly, in this approach, the genome was divided into 50 kb bins, and for each bin, nuclei were grouped into two bulks based solely on their genotype in that bin (MH63 or ZS97). Genome-wide differential CA analysis was then performed between the two bulks to test all ACRs simultaneously. By repeating this process across all genomic bins, a genome-wide association matrix was constructed, where each row represents an ACR and each column represents a genomic bin, with the *P* values along each row reflecting the genome-wide genotype-CA association profile of that ACR. An ACR with a significant *P* value at a particular bin is likely regulated by genetic variations within or near that bin, indicating a caQTL (Fig. 5a, Methods).

**Fig. 5:**
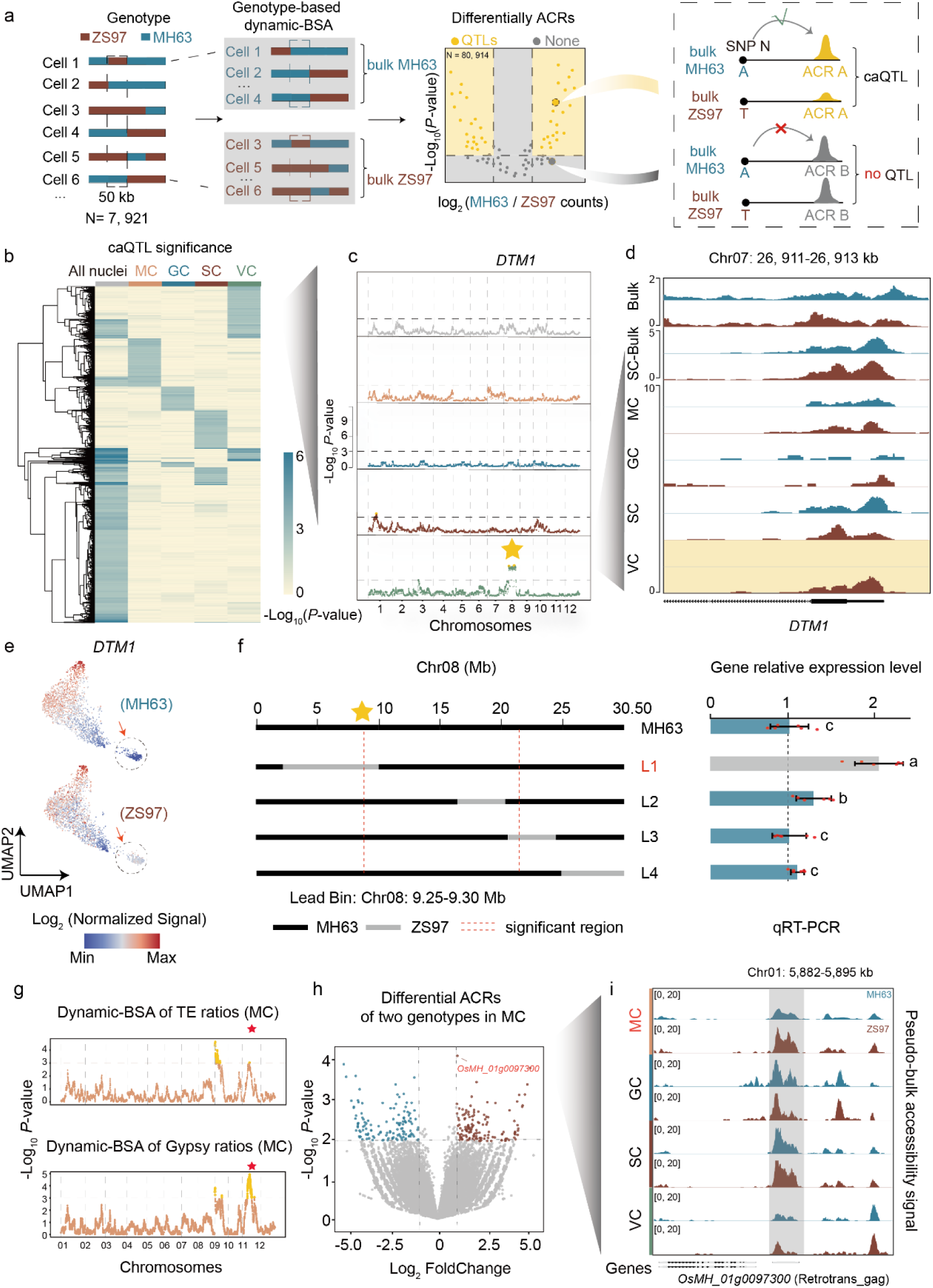
Dynamic-BSA maps the genetic basis of chromatin accessibility variation. **a**, Schematic of Dynamic-BSA. Left: Nuclei are assigned to MH63 or ZS97 bulks based on their genotype at each 50 kb genomic bin, followed by genome-wide differential chromatin accessibility (CA) analysis. Right: An accessible chromatin region (ACR) showing significant CA differences between genotypic bulks at a given bin is identified as a caQTL. **b**, Heatmap of caQTL significance across all nuclei and individual cell types. Rows represent caQTL loci, columns indicate cell populations, and color denotes −log_10_(*P*-value) from DESeq2; rows are hierarchically clustered. **c**, Dynamic-BSA plots for the *DTM1* promoter ACR (Chr07: 26,911-26,913 kb) across all nuclei and individual cell types. Yellow star, significant *trans*-caQTL detected in VC on chromosome 8. **d**, CA profiles at the *DTM1* promoter. Bulk, independent ATAC-seq data from MH63 and ZS97 pollen; Single Cell Bulk (SC-Bulk), pseudo-bulk of all single nuclei grouped by genotype; MC, GC, SC, and VC, cell-type-resolved pseudo-bulk tracks grouped by genotype at the chromosome 8 *trans*-caQTL. MH63 (blue) and ZS97 (brown) are shown. Note that the VC-specific CA difference is masked in the pooled nuclei data. **e**, UMAP projection colored by *DTM1* gene accessibility score, comparing nuclei with MH63 versus ZS97 genotypes at the chromosome 8 *trans*-caQTL. The dashed circle highlights the VC cluster. **f**, Validation using introgression lines. Left: Genotype schematics of four lines (L1-L4) with MH63 background carrying different ZS97 segments on chromosome 8 (Additional file 2: Table S4). Dashed lines indicate the extent of the significantly associated region. Right: qRT-PCR of *DTM1* expression (Additional file 2: Table S3). Different letters indicate significant differences (ANOVA, Tukey’s post-hoc analysis, *P* < 0.05). **g**, Dynamic-BSA of TE-derived read proportions in MC. Top: All TEs; bottom: *Gypsy*-type retrotransposons. Red stars indicate the most significant QTL for the *Gypsy* ratio on chromosome 11. **h**, Differential CA at chromosome 11 *Gypsy* QTL in MC. The most significantly differentially accessible locus, *OsMH_01g0097300* (encoding a Retrotrans_gag domain protein), is labeled. **i**, CA profiles of *OsMH_01g0097300* (Chr01: 5,882-5,895 kb) across cell types, grouped by genotype, showing MC-specific genotype-dependent accessibility.

We validated Dynamic-BSA using an introgression line R1, which carries a ZS97 genetic background with an introgressed MH63 segment on Chromosome 4 (17.8-20.7 Mb) (Fig. 5b, Additional file 1: Fig. S7a). Differentially accessible regions (DARs) between R1 and ZS97 showed highly significant overlap with ACRs identified by Dynamic-BSA as regulated by this introgressed interval (Fisher’s exact test, *P* value = 4.60 × 10^−14^ for the most significant 50 kb bin), providing strong cross-validation. Moreover, enrichment scores were substantially higher within the introgressed segment than in flanking regions, confirming the specificity of Dynamic-BSA (Additional file 1: Fig. S7a). Additionally, permutation tests in which nuclei were randomly assigned to bulks confirmed that genotype-based caQTLs were consistently more significant than those from random assignments (Additional file 1: Fig. S5b).

Using the single-nucleus data aligned to MH63RS3, we called 80,974 ACRs, each treated as a quantitative trait for genetic mapping (Methods). With 50 kb bins (7,921 bins genome-wide), Dynamic-BSA produced an 80,974 × 7,921 association matrix. Applying this framework to all 4,887 nuclei, we identified 8,051 significant caQTLs (FDR < 0.05, Additional file 2: Table S6, S7, Methods), comprising 1,747 *cis* (ACR-bin distance ≤ 5 Mb on the same chromosome) and 6,304 *trans* (Additional file 1: Fig. S6). To dissect cell-type-specific regulatory landscapes, we performed separate analyses for each cell population, revealing remarkable regulatory diversity: MC (170 *cis* and 2,457 *trans*), GC (186 *cis* and 1,372 *trans*), SC (370 *cis* and 2,601 *trans*), and VC (724 *cis* and 1,911 *trans*) (FDR < 0.05, Additional file 2: Table S6, S7). Combining the global and cell-type-stratified analyses yielded a total of 16,113 non-redundant significant caQTLs involving 14,319 ACRs. Of these, 48.6% (7,827: 896 *cis* / 6,931 *trans*) were cell-type-specific, whereas 39.3% (6,329: 1,222 *cis* / 5,107 *trans*) were detected only in the pooled analysis (Additional file 1: Table S7, Fig. 5b).

To illustrate the biological significance of cell-type-specific regulation, we examined an ACR (Chr07: 26,911-26,913kb) in the promoter of *DEFECTIVE TAPETUM AND MEIOCYTES 1* (*DTM1*) gene, a gene essential for early rice pollen development whose mutations impair pollen formation^37^. When analyzing all nuclei collectively, no caQTL was detected for this ACR, with no significant CA differences between MH63 and ZS97 genotypes (Fig. 5c). However, analysis of VC nuclei specifically revealed a *trans*-caQTL on chromosome 8 (9.05-22.5 Mb; the broad interval reflects suppressed recombination around the pericentromeric region). When we grouped VC nuclei based on their genotypes at this *trans*-caQTL, those with the ZS97 genotype showed significantly higher CA at the *DTM1* promoter compared to those with the MH63 genotype (Fig. 5d). Consistent with this chromatin state difference, UMAP visualization revealed markedly higher *DTM1* gene accessibility in ZS97-genotype VC nuclei compared to MH63-genotype VC nuclei (Fig. 5e).

To validate this regulatory relationship, we examined four introgression lines (L1-L4) with MH63 background containing different ZS97 segments on chromosome 8 (Additional file 1: Fig. S7c). qRT-PCR analysis revealed significantly higher *DTM1* expression in L1, which carries the ZS97 2.28-9.65 Mb segment overlapping with the associated region, compared to MH63 and the other lines (Fig. 5f). This experimental validation preliminarily confirms the regulatory relationship identified by Dynamic-BSA.

Beyond identifying *trans*-caQTLs, the cell-type-resolved chromatin maps generated by GAMETE also enable computational dissection of mechanistic basis of *cis*-caQTLs. While Dynamic-BSA maps *cis*-caQTL intervals, these regions typically harbor multiple linked variants, identifying the causal nucleotide requires modeling how each sequence change affects TF binding and CA. We addressed this using Basenji, a deep learning framework that predicts CA directly from DNA sequences, thereby capturing transcription factor binding affinities encoded within regulatory element^38^. Models trained on our cell-type-resolved MPS-ATAC-seq data mapped to the MH63 and ZS97 genomes achieved high accuracy (AUC = 0.939-0.976, Additional file 1: Fig. S8a, b). Using *in silico* mutagenesis, we interrogated a prominent SC-specific *cis*-caQTL at the promoter of *OsMH_05g0302900* (Additional file 1: Fig. S8c, d). The model pinpointed an A-to-G polymorphism between MH63 and ZS97 that was predicted to increase accessibility exclusively in SC, while models trained on VC, GC, and MC data predicted minimal effects (Additional file 1: Fig. S8e, f). This demonstrates that deep learning can complement Dynamic-BSA by resolving *cis*-caQTLs to candidate causal variants.

Collectively, these results demonstrate that Dynamic-BSA, complemented by deep learning-based sequence analysis, effectively overcomes the challenges of sparse single-nucleus data to reveal cell-type-specific genetic regulation of CA.

### Genetic determinants of TE accessibility revealed by Dynamic-BSA

Having established that *gypsy*-type retrotransposons exhibit distinctively elevated CA in MC (Fig. 2), we next investigated whether genetic variation contributes to the cell-type-specific differences in TE accessibility. Genome size variations among *Oryza* species are largely driven by differential TE expansion^20^, suggesting that the genetic control of TE activity varies between rice varieties. To map the loci underlying such variation, we extended our Dynamic-BSA framework by treating the proportion of TE-derived reads in individual nuclei as quantitative traits. We partitioned the genome into 50 kb bins (as illustrated in Fig. 4a) and, for each bin, classified all nuclei into two groups based on their genotype at that bin. Subsequently, we performed *t*-tests comparing proportions of TE-derived sequences between these two groups to identify genomic regions associated with TE accessibility (Methods).

The analysis revealed several significant QTLs associated with CA at TEs, with marked differences across both cell types and TE classes (Fig. 5g, Additional file 1: Fig. S9a). Notably, when restricting the analysis to *gypsy*-type retrotransposons in MC, we detected a major QTL on chromosome 11. This suggests that specific genetic variants on chromosome 11 exert a pronounced effect on *gypsy* retrotransposon accessibility in MC (Fig. 5g, Additional file 1: Fig. S9b-d). The distinct pattern of genetic control between TE classes suggests class-specific regulatory mechanisms operating in a cell-type-dependent manner.

To further dissect the genetic factors underlying this association, we performed differential CA analysis between MC nuclei grouped by their genotype at the *gypsy*-associated QTL on chromosome 11 (13.80-13.90 Mb). The most significantly differentially accessible locus was an ACR near *OsMH_01g0097300* on chromosome 1, encoding a putative TE protein containing the Retrotrans_gag domain, which displayed significantly higher accessibility in nuclei carrying the MH63 allele at the chromosome 11 QTL (fold change > 2, adjusted *P* value < 0.01; Fig. 5h-i), suggesting a *trans*-regulatory link between the chromosome 11 QTL and TE-related chromatin remodeling. Together, these results demonstrate that Dynamic-BSA can be extended from mapping regulatory variants at individual ACRs to dissecting the genetic architecture of aggregate chromatin phenotypes such as cell-type-specific TE accessibility.

## Discussion

GAMETE integrates high-throughput single-nucleus CA profiling with computational genotype inference and genetic mapping, enabling the simultaneous resolution of chromatin states, genotypes, and cellular identities from individual haploid nuclei. Applying this framework to hybrid rice pollen revealed distinct chromatin landscapes across cell types. Most notably, we discovered unexpectedly elevated CA at *gypsy* retrotransposons in MC, a phenomenon undetectable by standard pipelines that filter multi-mapping reads. Trajectory analysis linked this pattern to the distinctively low accessibility of RdDM components (*DCL3b*, *AGO4a*, *DRM2*) in MC, revealing a transient epigenetic surveillance gap between meiosis and the establishment of siRNA-mediated silencing in GC and SC^3,4^. This gap may allow retrotransposons to generate new insertions before being silenced^39^. Supporting this model, the two most accessible *gypsy* subfamilies in MC (Os0039_RIRE3 and Os0016) show the most pronounced copy number changes during rice domestication (Additional file 1: Fig. S3d). Given that *gypsy* expansion is the primary driver of genome size differences among *Oryza* species^40,41^, our findings provide a cellular and developmental context for understanding how transposon-associated genome evolution may arise, although the causal relationship between transient CA and long-term transposon expansion remains to be established. Consistent with this, key methylation and chromatin remodeling components (*OsDDM1b*, *SDG728*, *CMT3a*, *OsMET1b*) showed increased accessibility along the MC-to-GC trajectory but reduced accessibility toward VC, indicating coordinated reinforcement of epigenetic silencing in the germline^16,32^.

By exploiting the genetic basis of CA variation in F_1_ hybrid pollen, Dynamic-BSA identified 16,113 caQTLs, the vast majority exhibiting cell-type specificity. The *trans*-regulatory relationship between a chromosome 8 locus and *DTM1* accessibility, detected exclusively in VC and supported by expression analysis in introgression lines, exemplifies how cell-type-resolved analysis reveals regulatory mechanisms invisible to bulk approaches. Extending Dynamic-BSA to TE accessibility as a quantitative trait further identified a major QTL on chromosome 11 associated with *gypsy* accessibility in MC, providing a genetic entry point for the observed retrotransposon dynamics. Complementarily, a deep learning model trained on GAMETE data predicted CA from DNA sequence (AUC > 0.93), enabling *in silico* dissection of putative causal variants and functional elements underlying *cis*-caQTLs, thereby providing a starting point for nominating candidate upstream regulators.

Despite these advances, the sequencing depth inherent to single-cell data may affect the precision of cell-type annotation and caQTL detection, and integration with complementary modalities such as single-cell transcriptomics would provide a more comprehensive regulatory landscape. Direct cell-type-specific functional validation also remains technically challenging and will benefit from advances in single-cell manipulation technologies.

## Conclusion

We establish GAMETE as a broadly applicable framework for dissecting the genetic architecture of CA during gametogenesis. By overcoming the extreme data sparsity of single-cell profiling, GAMETE performs genetic mapping in haploid nuclei without the subsequent generations that conventional approaches require. Applied to rice pollen, it revealed a transient epigenetic surveillance gap that may contribute to *gypsy* retrotransposon expansion, identified thousands of cell-type-specific caQTLs, and traced SD loci to specific developmental stages, as demonstrated for a chromosome 4 locus detectable as early as MC. The framework could extend to any species in which haploid cells carrying natural genetic variation can be isolated. Moreover, its compatibility with index sorting, which records morphological parameters during fluorescence-activated sorting, potentially enables the integration of cellular phenotypes with genotypes and chromatin states at single-nucleus resolution.

## Methods

### Plant materials and growth conditions

SY63, MH63, ZS97 and their introgression lines were grown under normal agricultural conditions on the experimental farm of Huazhong Agricultural University, Wuhan, China. For single-cell analysis, SY63 pollen was harvested in two independent batches to generate two biological replicates for MPS-ATAC-seq library construction. For Dynamic-BSA validation, an introgression line (R1) was developed using ZS97 as the genetic background, with a specific segment from MH63 (Chromosome 4: 17.8-20.7 Mb) introgressed. For *DTM1* caQTL validation, four introgression lines (L1-L4) were developed using MH63 as the genetic background, carrying different ZS97 segments on chromosome 8 (Additional file 2: Table S4).

### Tissue preparation and nuclei extraction

The pollen was harvested as follows: Firstly, cut open rice spikelets, including the husks, and place them in pre-cooled 45% sucrose solution. The mixture was then pressed and stirred with a pestle for approximately 20 minutes to release the pollen, turning the solution yellow. Next, the yellow solution was filtered through a 140-mesh sieve to remove larger tissue chunks. The filtrate was collected using a 40 μm filter, trapping the pollen grains. The filter was rinsed with a small amount of 45% sucrose to obtain a concentrated pollen solution, which was then collected into a 1.5 mL centrifuge tube and gently centrifuged for about 5 seconds. Then, remove the supernatant, add twice the volume of water to dilute the sucrose concentration to 15%, mix at room temperature, vortex every five minutes to assist pollen grain expansion and rupture. After 20 minutes, the solution was centrifuged at 500 ×g for 10 minutes at 4°C, and the supernatant was discarded. An equal volume of chopping buffer (without Triton X-100) was added, and the mixture was pipetted several times to ensure thorough mixing. Subsequently, the solution was filtered through a 30 μm filter, and approximately 1 mL of filtrate was collected. To stain the nuclei, 15 μL of 0.1 mg/mL DAPI was added and incubated for 5 minutes. Finally, the stained pollen nuclei were sorted by flow cytometry, with those showing haploid DNA content selected.

### Tn5 production and transposome assembly

Tn5 production and transposome preparation were performed as previously described^42^. All the oligonucleotides were synthesized by Sangon Biotech (Additional file 2: Table S1, S2). For MPS-ATAC-seq, ME-S1 to S24 (each containing a unique barcode), and ME-B were annealed to MErev separately to form double-stranded MEDS-S and MEDS-B complexes. Then, a 1:1 ratio of MEDS-Ss and MEDS-B was assembled with in-house Tn5 at room temperature for 1 hour to form the final 24 transposomes. For ATAC-seq, ME-A was used instead of ME-S for the assembly. The transposomes were stored at −20 °C.

### MPS-SC-ATAC-seq library preparation

The process is briefly described as follows. First, 30,000 pollen nuclei per tube were sorted into a total of 24 tubes, each tagmented with a Tn5 enzyme containing one of 24 unique barcodes. The tagmentation reaction was carried out by adding 0.5 μL Tn5-MEDS-S/B complex, 10 μL 5′ reaction buffer, and 29 μL ddH₂O into each of the 24 tubes, followed by incubation at 37°C for 30 minutes. Next, the reaction was terminated at the completion of tagmentation, and a second round of sorting was performed. Nuclei carrying different barcodes were sorted tube by tube into a 384-well plate pre-loaded with 24×16 primer combinations (Additional file 2: Table S1), and immediately stored at −80°C, where the process could be paused. Subsequently, the 384-well plate was taken out of the refrigerator, quickly thawed, and nuclei were lysed using proteinase K at 50°C for 40 minutes, followed by inactivation at 70°C for 20 minutes. NEBNext® High-Fidelity 2× PCR Master Mix (NEB, M0541) was added to each well for PCR amplification. The amplification conditions were as follows: 72°C for 5 minutes; 98°C for 2 minutes; then 16 cycles of 98°C for 10 s, 59°C for 30 s, and 72°C for 50 s; followed by a final extension at 72°C for 5 minutes, and held at 4°C. The PCR products from all 384 wells were pooled and purified using the Takara Mini DNA Purification Kit (Takara, 9761) with a QIAvac vacuum system. The eluted DNA was immediately subjected to 1.5× AMPure beads (Beckman, A63881) purification and eluted with 20 μL ddH₂O. The bead-eluted DNA was then subjected to a second round of PCR amplification with Ad1.sc.6/7 and P7 primers and NEBNext® High-Fidelity 2× PCR Master Mix. The amplification conditions were as follows: 98°C for 2 minutes; then 8 cycles of 98°C for 10 s, 63°C for 30 s, and 72°C for 50 s; followed by 72°C for 5 minutes, and held at 4°C. Finally, the library was purified again with 1.5× AMPure beads and eluted with 20 μL ddH₂O prior to DNA quantification. PE150 sequencing was performed on an Illumina NovaSeq 6000 with a sequencing depth of 150 Gb per sample.

### MPS-ATAC-seq data splitting

Barcodes are essential for distinguishing single cell data. The raw sequencing data was organized into 48 FASTQ files (24 for READ1 and 24 for READ2), each corresponding to one of the 24 i7 index sequences. For each i7-indexed file, a combination of X_i5_ and Y_Tn5 libraries_ generates X×Y barcode sequences (specifically, 16×24). The first 35 bases of READ1 are designated as the barcode for data splitting (detailed barcodes are listed in Additional file 2: Table S1, S2). The 384 most frequent (16×24) barcode combinations across all READ1 sequences were extracted and assessed to ensure they matched with the barcodes derived from the sequencing libraries. In the preprocessing of raw FASTQ files, a two-stage splitting strategy was applied. Initially, the READ1 file was split into 384 separate FASTQ files, each corresponding to one of the 384 barcodes. Subsequently, leveraging the information from READ2, UMItools^43^ was employed to further delineate these files into separate READ1 and READ2 files for each of the 384 single cells. Consequently, each raw FASTQ file was resolved into 384 distinct single-cell files. Applying this procedure to each batch of MPS-ATAC-seq data generated a comprehensive dataset of 9,216 single-cell files (24 × 384 cells per batch). The detailed workflow is as follows: First, the *fastx_barcode_splitter.pl* tool (https://github.com/lianos/fastx-toolkit/) was used to split the raw READ1 file into 384 single-cell datasets based on the barcode combinations, allowing for one mismatch with the parameter “--mismatches 1”. This step produced 24 × 384 READ1 datasets. Next, *repair.sh* from BBMap (https://github.com/BioInfoTools/BBMap) was used to extract READ2 sequences from every READ1 file generated in the previous steps. Finally, a total of 24 × 384 READ1 and 24 × 384 READ2 FASTQ files were obtained.

### MPS-ATAC-seq data processing

After splitting the single-cell data, reads were mapped to three reference genomes using BWA-MEM^44^: the Nipponbare genome (MSUv7) was used for cell clustering, chromatin accessibility quantification, and TE analysis, as it provides the most comprehensive functional annotation among available rice genomes; the two parental genomes MH63RS3 and ZS97RS3^35^ were used for genotype inference and allele-specific analyses. Specifically, the MH63RS3 genome was used for CA quantification to support the subsequent Dynamic-BSA analysis. Mapped reads with a MAPQ score below 30, PCR duplicates, and mapping to mitochondrial and chloroplast genome were filtered using SAMtools^19^ to ensure high-quality aligned data. BAM files were then converted to BED format^45^ and Tn5 insertion sites were adjusted according to strand orientation: the forward strand shifted by +4 bp, while the reverse strand shifted by −5 bp. To ensure high data quality, we applied a hierarchical filtering strategy based on genomic coverage uniformity and genotype consistency. First, to exclude nuclei that may have ruptured or fragmented during isolation (leading to partial genomic data loss), we compared read distribution of single nuclei against bulk ATAC-seq data from the same tissue (SY63 pollen). The genome was divided into 5 Mb bins, and a chi-square test was performed to assess the discrepancy in coverage distribution between single-cell and bulk data. Nuclei showing significant deviation were considered incomplete and excluded. Second, to filter out potential contamination or mixed cells, we applied an HMM to correct genotypes (see Methods: “Genotype correction by hidden Markov model”). We calculated the genotype consistency before and after HMM correction. Nuclei with low consistency were discarded, as high correction rates imply potential contamination. Based on these metrics, we retained 10,205 nuclei (genotype consistency ≥ 0.76, chi-square deviation score ≤ 10, and Tn5 insertions ≥ 1,000) for peak calling in the Dynamic-BSA analysis (Additional file 1: Fig. S1f). For downstream genotyping analysis, which requires higher precision, we applied a stricter threshold of genotype consistency ≥ 0.79. This yielded 5,190 high-quality nuclei, which were further processed using ArchR for standard quality control (see Methods: “Cell clustering and annotation”) and outlier removal, resulting in a final dataset of 4,887 nuclei. For TE-related analyses, a parallel set of BAM files was generated without MAPQ filtering (e.g., MAPQ ≥ 0). Because TEs typically exist as multiple genomic copies, standard MAPQ thresholds (e.g., MAPQ ≥ 30) disproportionately exclude reads derived from repetitive elements, leading to underestimation of TE accessibility. The unfiltered BAM files were used for all TE quantification, visualization, and TE-related QTL analyses described below, unless otherwise specified.

### Cell clustering and annotation

A custom ArchRGenome^15^ for rice was constructed using genomic sequences from the *BSgenome.Osativa.MSU.MSU7* R package and gene annotation from RAP-DB (https://rapdb.dna.affrc.go.jp/). Arrow files were generated with the following filters: minTSS = 2, bcTag = “RG”, minFrags = 500, maxFrags = 10000, promoterRegion = c(3000, 100), excludeChr = c(“chrUn”, “chrSy”). An ArchRProject was created with default parameters. Dimensionality reduction was performed using LSI with iterations =2, varFeatures = 22000, and dimsToUse = 1:30. Gene activity scores were derived using the addGeneScoreMatrix function. Cell types were annotated based on established marker genes from rice pollen develop mental studies^23,24,46,47,48,49,50^, selected by cross-referencing their expression specificity in publicly available microarray datasets^51,52^ and confirmed by cell-type-preferential CA patterns in our single-nucleus data (Fig. 1c,d, Additional file 1: Fig. S2b, g). Specifically, MC was identified by *CRC1*^46^ and *ZEP1*^47^; GC by *OsGCS1*^50^ and *OsDUO1*^21^; SC by *OsGEX2* and *Os08g0459100*^24^; and VC by *VCS1*^23^ and *OsMYB80*^48,49^.

### Identification of accessible chromatin regions

In this study, three distinct sets of ACRs were generated for quality control, genetic analysis, and cell-type characterization, respectively. 1, Reference set for quality control: To assess single-nucleus data quality (e.g., FRiP calculation), we utilized an independent SY63 pollen bulk ATAC-seq dataset. Reads were mapped to the Nipponbare reference genome (MSUv7), yielding 53,619 ACRs. Peak calling was performed using MACS2^53^ with parameters optimized for ATAC-seq: “-g 3.0e8 -nomodel -extsize 38 -shift -15 -keep-dup all -B -SPMR -call-summits”. 2, Set for Dynamic-BSA analysis: For genetic mapping and Dynamic-BSA, MPS-ATAC-seq data from filtered nuclei were merged and mapped to the MH63RS3 reference genome. This process yielded a comprehensive set of 80,974 ACRs, identified using MACS2 with the same parameters as above. 3, Set for cell-type-preferential ACRs analysis: To identify ACRs with cell-type-preferential accessibility, we generated a reproducible peak set using the “addReproduciblePeakSet” function in ArchR by default parameters. This resulted in 55,044 high-confidence ACRs. Based on this set, we further identified cell-type-preferential patterns using a maximum gap analysis. To identify ACRs with cell-type-preferential accessibility patterns, single-cell data were aggregated by cell type to generate pseudo-bulk profiles for MC, GC, SC, and VC. For each of the 55,044 ACRs, normalized accessibility was calculated in each pseudo-bulk profile and log_2_-transformed. The four values were then sorted in ascending order, and the maximum gap between consecutive values was computed to quantify cell-type-specific variation. ACRs were retained for pattern analysis if they satisfied two criteria: **(1)** the maximum accessibility value exceeded the genome-wide median, and **(2)** the maximum gap exceeded log_2_(3), corresponding to at least a 3-fold difference between adjacent cell types.

### RNA-seq datasets processing and accessibility-expression comparison

Publicly available rice pollen RNA-seq datasets (GSE235680 and GSE50777) were downloaded from NCBI GEO^16,17^. Transcript abundance was quantified using Salmon^54^ against the IRGSP-1.0 reference transcriptome. TPM values were used for downstream visualization and comparison. For CA, each gene was represented by its single most-accessible peak within a strand-aware window from 3 kb upstream to 1 kb downstream of the TSS, defined as the peak with the highest CPM across the four MPS-ATAC-seq pseudobulks; the per-cell-type value was the CPM of that peak in each pseudobulk. To evaluate the relationship between CA and gene expression, cell-type specificity scores were calculated independently for MPS-ATAC-seq and public RNA-seq data. For each gene, specificity was defined as:

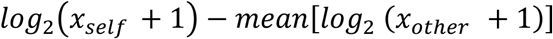

where *x_self_* represents the accessibility (CPM) or expression (TPM) in the focal cell type and *x_other_* represents the corresponding values in the remaining cell types. For RNA-seq data, specificity scores were calculated separately within each study to minimize systematic differences caused by distinct library preparation protocols. To enrich informative loci, a focused gene set was generated by taking the union of the top 500 cell-type-enriched genes from each MPS-ATAC-seq and RNA-seq category, yielding 3,262 genes shared between accessibility and expression datasets for downstream correlation analysis. Pearson and Spearman correlations were subsequently calculated using the union of cell-type-enriched genes identified from both datasets.

### Identification of stamen-preferentially expressed genes

RNA-seq data from multiple tissues spanning the full life cycle of rice were downloaded under the accession number PRJNA940508^55^. To identify genes preferentially expressed in stamen, differential expression analysis was performed using DESeq2, comparing stamen against all other tissues pooled together. Genes with a fold change ≥ 2 and an adjusted *P* value ≤ 0.05 were defined as stamen-preferentially expressed genes, yielding 2,586 genes. These genes were used to evaluate cell-type-specific CA patterns at functionally relevant loci by deepTools^56^ “computeMatrix scale-regions” function (Fig. 1h).

### Pseudo-time analysis

To analyze differentiation trajectories across different cell types, pseudotime analysis was performed using Monocle 2^57^. The CellDataSet object was constructed using the PeakMatrix generated by ArchR. ACRs detected in only a small number of cells were filtered out. DARs were then identified using the differentialGeneTest function with the parameter fullModelFormulaStr = “∼Clusters”. Dimensionality reduction was performed using the reduceDimension function with the method “DDRTree”, enabling visualization and interpretation of the two-dimensional cellular trajectory. Finally, the orderCells function was applied to arrange cells along the pseudotime trajectory, thereby reconstructing the differentiation process. Adopting a strategy similar to Cicero^58^, all single cells were divided into ten groups along the pseudotime trajectory, with each group representing a distinct segment of pseudotime and effectively partitioning the continuum into ten equal bins. Dynamic changes of pseudotime-specific ACRs across these groups were then analyzed, allowing characterization of CA patterns and identification of temporal regulation throughout the differentiation process.

### Identification of *cmt3a* DMRs

BS-seq data of rice sperm cells was downloaded under the accession number GSE235680^16^. To identify differentially methylated regions (DMRs), the rice genome (MSUv7) was divided into 100-bp bins. Bins exhibiting a CHG methylation difference of at least 0.3 between the wild-type and *cmt3a* mutant sample were defined as DMRs. Finally, overlapping DMRs (with at least a 1-bp overlap) were merged.

### Genotype determination in single pollen nuclei

To determine genotypes from single-cell ATAC-seq data of the F_1_ hybrid rice SY63 (ZS97 × MH63), we developed a multi-step computational pipeline. First, sequencing reads were independently aligned to both parental reference genomes (MH63RS3 and ZS97RS3) using BWA. For each read, alignment scores (AS) were compared between the two genomes to determine its parental origin. Reads with identical AS for both genomes were discarded as non-informative, while those with differential scores were retained for genotype inference. To overcome the inherent sparsity and noise of single-cell data, genotype information was aggregated across non-overlapping 10 kb genomic windows. Within each window, a genotype call was made only when > 80% of informative reads consistently supported one parental origin; otherwise, the region was marked as undetermined. We further filtered potential alignment artifacts by excluding reads that mapped to different chromosomes or exhibited positional discrepancies exceeding 5 Mb between the two reference genomes.

### Genotype correction by hidden Markov model (HMM)

To refine the initial genotype calls and account for the continuous nature of recombination blocks, we implemented an HMM adapted from our previous work^8^. This approach leverages the principle that adjacent genomic regions likely share identical genotypes due to limited recombination events during meiosis. In the HMM framework, hidden states represent the true parental origin (MH63 or ZS97), while observed states correspond to the initial genotype calls in each window. The emission probability was set to 0.99 (match) and 0.01 (mismatch), reflecting an assumed sequencing error rate of 1%. For undetermined windows, the emission probability was set to 0.5 for each genotype. Transition probabilities were modeled as functions of physical distance between adjacent genotyped segments, reflecting recombination probability. For two segments separated by distance d (kb), the transition probability was calculated by converting physical distance to genetic distance using the average recombination rate in rice (250 kb/cM), then applying Haldane’s mapping function to obtain recombination fraction r:

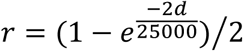

The transition probability matrix is shown in Fig. 4a. The model was initialized with equal parental genotype probabilities (0.5:0.5) and solved for each chromosome independently using the Viterbi algorithm. After HMM correction, adjacent segments with identical inferred genotypes were merged into continuous recombination blocks, while segments with ambiguous calls were designated as missing data. Recombination breakpoints were defined as the midpoints between segments where genotype transitions occurred.

### Identification of segregation distortion loci

SD occurs when observed genotype frequencies deviate significantly from the expected Mendelian ratios. In this study, we identified these events by examining the distribution of ZS97 and MH63 genotypes within genomic bins (10 kb window). After constructing the genotype map, the number of cells with each genotype was counted for every bin. A Chi-square test was performed for each bin to assess deviation from the expected 1:1 segregation ratio. Given that the 10 kb window size might introduce stochastic noise, we calculated the mean *P* value across a 100 kb window (averaging ten 10 kb bins). Ultimately, genomic regions with a mean *P* value < 0.001 were identified as SD loci.

### Comparative analysis of recombination landscapes

We quantified recombination intensity across the genome for both the single-nucleus pollen population generated in this study (mapped to MH63RS3) and a previously published MH63 × ZS97 recombinant inbred line population (n = 210, mapped to MSUv6)^33^. To facilitate direct comparison between the two reference genomes, we performed coordinate conversion (liftover) using 50 kb windows to establish homologous genomic intervals. To ensure statistical robustness, recombination rates were calculated using a sliding window approach with a 5 Mb window size and a 2.5 Mb step size. For the chromatin analysis, the genome was partitioned into non-overlapping 2.5-Mb windows, which were ranked by recombination rate in the pollen population; the top 10% were defined as high-recombination regions and the remainder as background. For each window, chromatin accessibility was quantified from merged MPS-ATAC-seq reads (all four cell types) as log₂(CPM) and compared between the two groups using a two-sided *Wilcoxon* rank-sum test.

### Dynamic-BSA mapping of caQTLs

The MH63RS3 genome was partitioned into 7,921 non-overlapping 50 kb bins. For each bin, haploid nuclei were assigned to MH63, ZS97, or undetermined genotype based on the inferred parental origin in that interval. Nuclei with undetermined calls were excluded, and the remaining nuclei were grouped into MH63 and ZS97 genotype bulks. Subsequently, to enable dispersion estimation for differential testing, the nuclei within each genotype group were randomly split into four pseudo-replicates. For each nucleus, accessibility at each ACR was binarized (presence or absence of Tn5 insertion), and the binary signals were then aggregated within each replicate to generate an ACR-by-sample count matrix (80,974 ACRs × 8 pseudo-bulk samples). Differential accessibility between the MH63 and ZS97 bulks was then tested on these aggregated counts using DESeq2 (default settings). Repeating this procedure for all bins generated a genome-wide association matrix of *P* values (80,974 ACRs × 7,921 bins).

### Permutation-derived filtering of candidate caQTLs

To increase stringency while accounting for ACR-specific variability, we performed 10 label permutations by shuffling nucleus-to-genotype assignments and reran the complete Dynamic-BSA pipeline for each permuted dataset. In the observed data, ACR-bin associations with nominal *P* < 0.005 were first retained. To reduce redundancy caused by linkage, only the most significant association per chromosome was retained for each ACR. For each permutation and each ACR, we recorded the minimum *P* value across the genome to define an ACR-specific null extreme. An observed ACR-bin association was retained as a high-confidence caQTL only if its nominal *P* value was more significant than all 10 permutation-derived minima for that ACR.

### *cis*/*trans* classification and FDR estimation

Significant caQTLs were classified as *cis* if the associated bin was within 5 Mb of the ACR on the same chromosome, and as *trans* otherwise. To assess the empirical false discovery burden, we calculated permutation-based FDR separately for *cis* and *trans* signals as

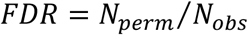

where *N*_obs_ is the number of retained high-confidence ACR-bin associations in the observed dataset after applying both the nominal cutoff and the permutation-derived extreme-value filter, and *N*_perm_ is the corresponding number in the permuted datasets after applying the same procedure. Overall FDR was calculated analogously. The entire analytical pipeline was also performed independently within each annotated cell type to characterize cell-type-specific genetic effects.

### Genotype validation of introgression line

To validate the genotype of the ZS97 introgression line R1, we performed ATAC-seq and aligned the sequencing reads to the MH63RS3 and ZS97RS3 reference genomes, respectively. Genotype validation was conducted by applying the identification and correction procedures described in the “Genotype determination in single pollen nuclei” and “Genotype correction by hidden Markov model” sections. Similarly, RNA-seq data were generated for the four MH63 introgression lines (L1-L4), and their genotypes were confirmed using the identical workflow.

### qRT-PCR validation of *DTM1* expression

Anthers were collected from MH63 and the four introgression lines (L1-L4) at the heading stage, with at least three biological replicates per line. Total RNA was extracted using the RNAprep Pure Micro Kit (TIANGEN, DP420) following the manufacturer’s protocol. First-strand cDNA was synthesized using the HiScript IV 1_st_ Strand cDNA Synthesis Kit (Vazyme, R412). qRT-PCR was subsequently performed using Taq Pro Universal SYBR qPCR Master Mix (Vazyme, Q712) on an Applied Biosystems^®^ QuantStudio^TM^ 7 Flex Real-Time PCR System. Rice Ubq (*LOC_Os03g13170*) was used as the internal control to analyze *DTM1* expression levels with the comparative critical threshold (2^−ΔΔCT^) method. Primers used for qRT-PCR are listed in Additional file 2: Table S3. Differences among groups were evaluated using one-way analysis of variance (ANOVA) followed by Tukey’s Honest Significant Difference (HSD) post-hoc test. A *P*< 0.05 was considered statistically significant.

### Quantification of cell-type-specific TE accessibility

To characterize TE accessibility profiles across different cell types, we utilized the rice TE annotation generated by the Extensive de-novo TE Annotator (EDTA) pipeline, based on the MSUv7 reference genome^18^. To enhance annotation reliability, TEs shorter than 50 bp were excluded. Two MAPQ filtering strategies were applied to evaluate the impact of multi-mapping reads on TE accessibility quantification. For each single nucleus, TE accessibility was quantified as the proportion of reads overlapping with annotated TEs relative to total mapped reads. This metric was calculated under both MAPQ ≥ 30 (standard filtering, retaining only uniquely mapped reads) and MAPQ ≥ 0 (no filtering, retaining all aligned reads including multi-mapping reads). Comparative analysis revealed that the MAPQ ≥ 30 filter discarded substantial genuine signals from repetitive elements, particularly in MC (Fig. 2a). Therefore, all subsequent TE analyses were performed using unfiltered data (MAPQ ≥ 0). To identify TE subfamilies exhibiting accessibility levels beyond the expectation from their genomic abundance, we calculated the total genomic length occupied by each TE subfamily and computed correlation coefficients between genomic length and the proportion of accessible reads for each cell type separately (Additional file 1: Fig. S3a).

### Identification of cell-type-specific TE regulatory loci

To identify genomic regions influencing TE accessibility across cell types, we employed the Dynamic-BSA framework using the unfiltered BAM files (MAPQ ≥ 0). The genome was first partitioned into 50 kb non-overlapping bins. For each bin, single cells were stratified into MH63 and ZS97 genotype groups using the previously established genotype maps, and the proportion of TE-derived reads (relative to total mapped reads per nucleus) was calculated for each group. We then performed *t*-tests to compare the TE proportions between the two groups, identifying statistically significant (*P* value < 0.001, Fig. 5g) regions that potentially regulate TE accessibility in a genotype-dependent manner.

### Deep learning model construction

We utilized the Basenji deep learning framework, which we previously optimized for modeling CA in the rice genome^38,55^; similar parameters were used in this study. Briefly, sequencing reads from individual cell were independently mapped to the MH63 and ZS97 genomes using BWA. Reads with a mapping quality score below 30, PCR duplicates, and reads mapping to mitochondrial or chloroplast genomes were filtered out using SAMtools. Subsequently, single-cell BAM files corresponding to identical cell types were merged and converted into BigWig files using the *bam_cov.py* script^38^. To prepare input files for the deep learning model, we employed the *basenji_data.py* script with the following parameters: “-l 16384 -p 20 -t Chr05 -v.1”. Chromosome 5 was used as the test set, and 10% of the remaining data (with random seed set to 1) was used as the validation set. The model was trained on an NVIDIA Tesla P100 GPU using the *basenji_train.py* script with the parameters: “--augment_rc --ensemble_rc --augment_shifts ‘1, 0, −1’”. The batch buffer parameter was set to 0. To evaluate the impact of sequence variations within ACRs, we performed in silico mutagenesis using the *basenji_sat_bed.py* script. The effect of each variation was quantified by calculating the signal difference between the reference and altered sequences. To predict the CA of orthologous ACRs between the two genomes, sequences were centered on each ACR and extended to a total length of 32,768 bp. Sequences exceeding the boundaries of the corresponding chromosome were excluded. Finally, sequences were extracted using the BEDTools getfasta command^45^. Furthermore, the *basenji-predict_bed.py* script was modified to accept FASTA files as input.

## Data availability

Sequencing data generated in this study are available in the Genome Sequence Archive (GSA) of the China National Center for Bioinformation (CNCB) under the BioProject ID: PRJCA058132 (GSA accession: CRA038772). The code used in this study is available upon reasonable request.

## Acknowledgments

The computations in this paper were run on the bioinformatics computing platform of the National Key Laboratory of Crop Genetic Improvement, Huazhong Agricultural University. This work was supported by grants from the National Natural Science Foundation of China (U23A20189), the Biological Breeding-National Science and Technology Major Project (2023ZD04076), the Hubei Provincial Natural Science Foundation of China (2023AFA043), the Earmarked Fund for China Agriculture Research System (CARS-01), and the Fundamental Research Funds for the Central Universities (2662023PY002).

## Contributions

W.X. designed and supervised this study. Y.L, Junjie.L, H.L and M.L analyzed the data. Y.L, Junjie.L, C.X and W.X wrote the manuscript. Y.L and W.X revised the manuscript. C.X, Junjie.L, J.Y, H.Y and T.Z performed the experiments. Z.Z, Jiacheng.L, L.M and X.H provided ideas for data analysis. Y.X, S.Y and Y.O participated in experiment design and discussions. All the authors read and approved the manuscript.

## Ethics declarations

### Competing interests

The authors declare no competing interests.

## Notes

### Competing Interest Statement

The authors have declared no competing interest.

