## Supplementary Figures for "GAMETE maps the genetic architecture of chromatin accessibility in rice pollen at single-nucleus resolution"

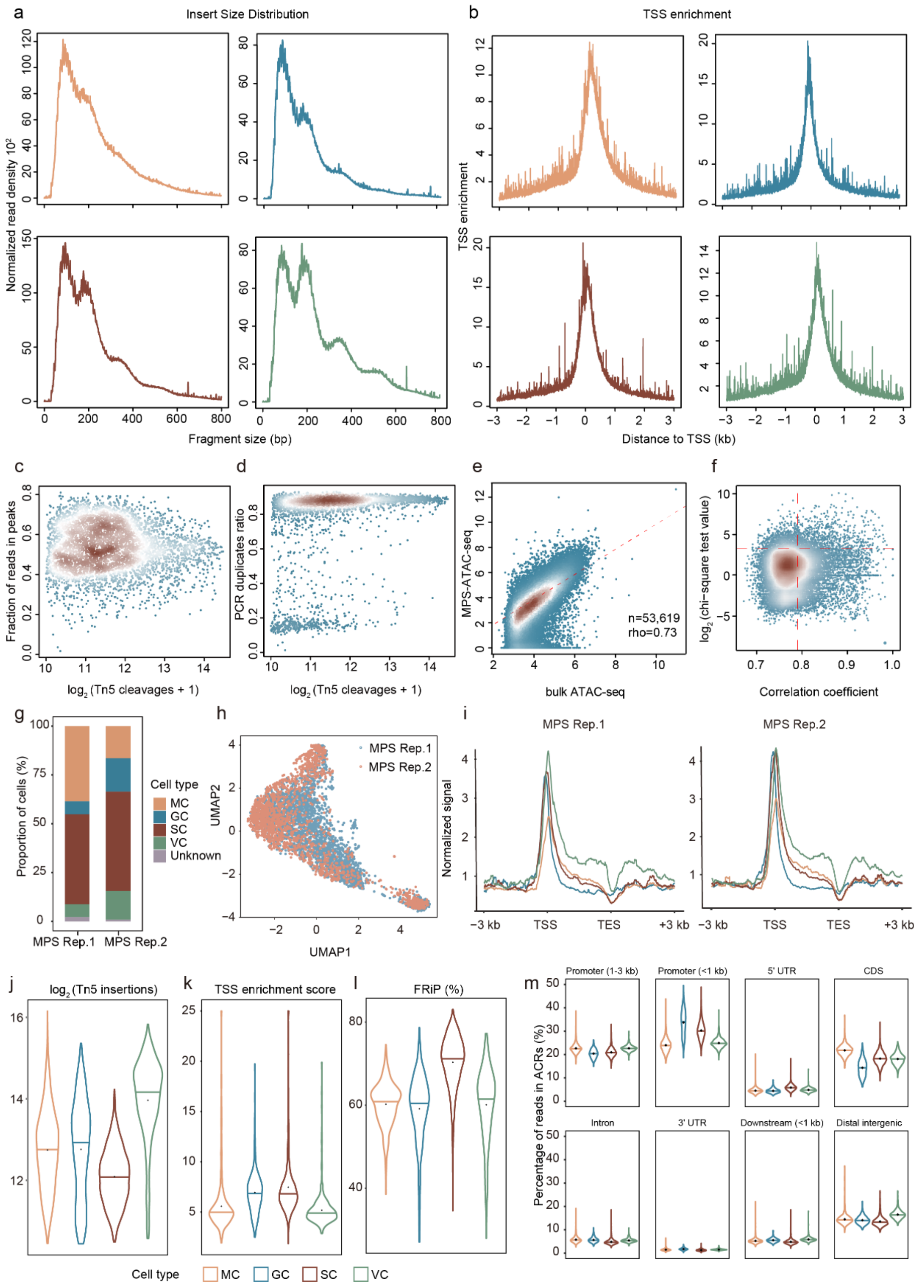


**Fig S1. Overview and quality control of MPS-ATAC-seq data. a**, Fragment size distributions of MPS-ATAC-seq reads across different cell types. **b**, Transcription start site (TSS) enrichment profiles across different cell types. The enrichment scores were calculated using Tn5 insertions in flanking regions (-3 kb to -2.5 kb and 2.5 kb to 3 kb) as background. **c-d**, Density scatterplots showing sequencing depth ($log_2$-transformed Tn5 insertion counts) versus the fraction of reads in peaks (FRiP) (**c**) and PCR duplicate rates (**d**). **e**, Correlation of normalized chromatin accessibility signals at the union of peaks between bulk ATAC-seq and aggregated MPS-ATAC-seq data (n = 53,619, Spearman's $\rho$ = 0.73). **f**, Quality control filtering of single cells based on their correlation of genotype consistency (x-axis) and log$_2$(chi-square test value) (y-axis). Red dashed lines indicate the thresholds used to exclude cells with aberrant chromatin profiles. **g**, Cell type composition across biological replicates. **h**, UMAP projection of single nuclei colored by replicate. **i**, Distribution of fragments within 3 kb upstream and downstream of stamen-specific genes across different cell types in two replicates. **j-l**, Violin plots comparing quality metrics across the four cell-types: log$_2$(Tn5 insertions) **(j)**, TSS enrichment scores **(k)**, and FRiP scores **(l)**. The horizontal line represents the median values, and the black dot represents the mean values. **m**, Genomic distribution of ATAC-seq reads across eight distinct genomic annotations features. Genomic regions are categorized as follows based on their positions: promoter 1-3 kb (1 to 3 kb upstream of promoters), promoter <1 kb (within 1 kb upstream of promoters), 5′ UTR (5′ untranslated region), CDS, intron, 3′ UTR (3′ untranslated region), downstream < 1 kb (within 1 kb downstream of genes), and distal intergenic (encompassing all remaining genomic regions not classified above).

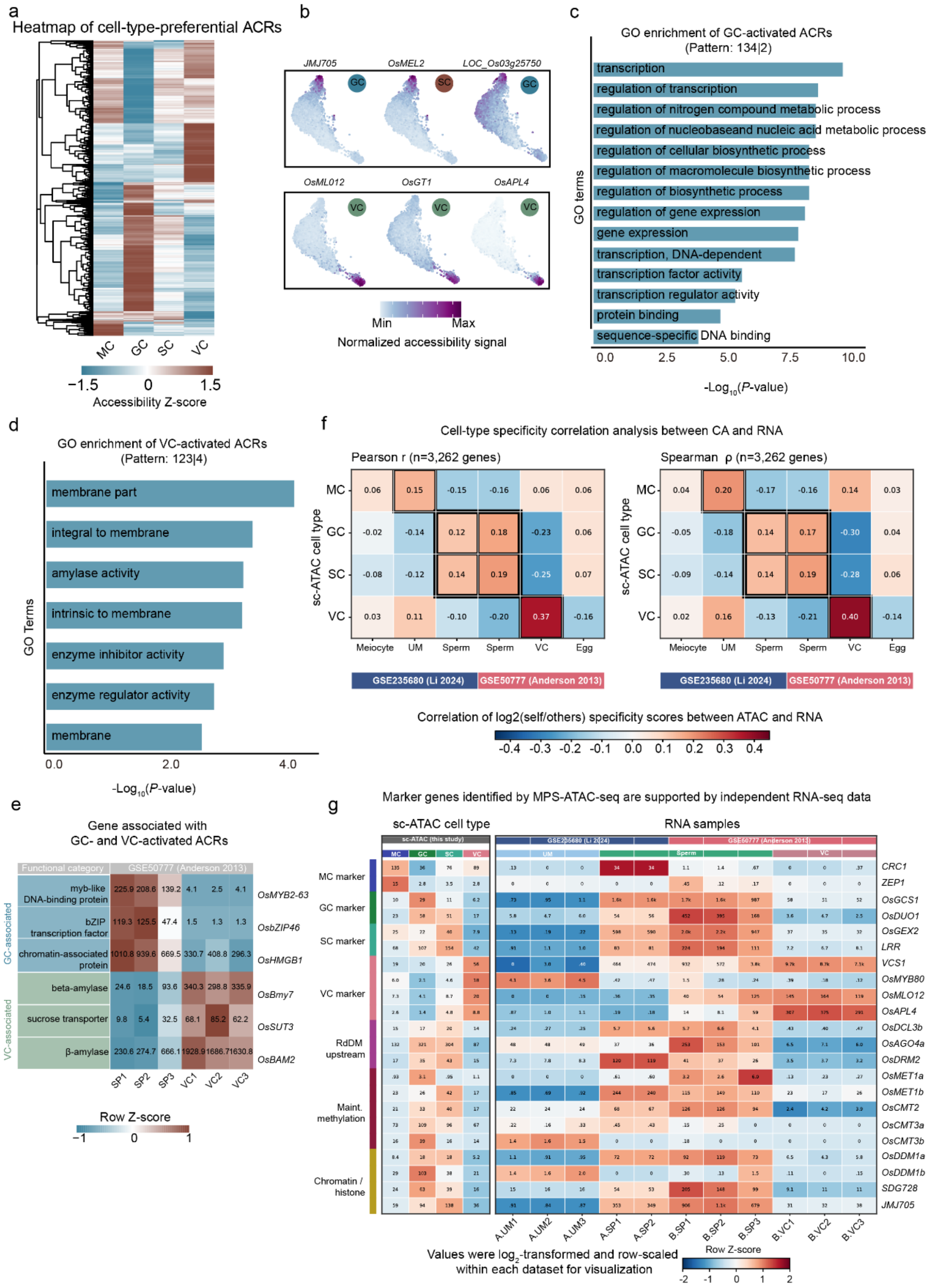


**Fig S2. Cellular heterogeneity revealed by MPS-ATAC-seq data. a,** Heatmap displaying the chromatin accessibility of cell-type-preferential ACRs across different cell types. **b,** Chromatin accessibility signal at regulatory regions of cell-type-specific marker genes. **c,d,** GO enrichment analysis of GC-preferential (pattern: 134|2) and VC-preferential ACRs (pattern: 123|4). Pseudo-bulk profiles were generated for MC (denoted as 1), GC (2), SC (3), and VC (4). Patterns represent accessibility levels sorted in ascending order, where the vertical bar (|) marks the maximum gap between consecutive values. For example, 134|2 indicates that the accessibility of cell type 2 (GC, right of the gap) is significantly higher than that of types 1, 3, and 4. **e**, Heatmap showing transcript abundance of six representative genes drawn from functional categories enriched among genes associated with GC- and VC-activated ACRs. Expression is from the rice pollen RNA-seq dataset (Anderson et al., 2013; three sperm-cell replicates, SP1-SP3; three vegetative-cell replicates, VC1-VC3). Cell colors represent row-scaled Z-scores of $log_2$-transformed TPM values, whereas numbers indicate the corresponding raw TPM values. **f**, Pearson correlation coefficients and Spearman rank correlations between CA specificity scores (Methods) derived from four MPS-ATAC-seq cell-types (MC, GC, SC and VC; rows) and expression specificity scores derived from six gametophytic cell-type groups (columns), each averaged over its biological replicates, across two independent transcriptome datasets (GSE235680, blue; GSE50777, red). **g**, Heatmaps showing promoter CA (left) and transcript abundance (right) for 22 representative marker and regulatory genes. Genes are grouped into seven functional categories, including cell-type marker genes, RdDM-related genes, maintenance DNA methylation genes, and chromatin/histone regulators. For the CA panel, each gene is represented by its single most-accessible peak within a strand-aware [TSS − 3 kb, TSS + 1 kb] window. The selected peak is the one with the highest CPM in any of the four sc-ATAC pseudobulks (MC, GC, SC, VC), and the per-cell-type values shown are the CPM of that peak in each of the four pseudobulks. For the expression panel, values represent TPM-normalized RNA abundance from two rice pollen transcriptome datasets: GSE235680 (Li et al., 2024; UM and sperm samples) and GSE50777 (Anderson et al., 2013; sperm and vegetative-cell samples). Colored bars above the RNA heatmap indicate dataset origin and cell type. Cell colors indicate row-scaled Z-scores calculated from $log_2$-transformed values within each dataset, whereas numbers shown in each cell represent the corresponding raw CPM (accessibility) or TPM (expression) values. UM: unicellular microspore; SP: sperm cell; VC: vegetative cell.

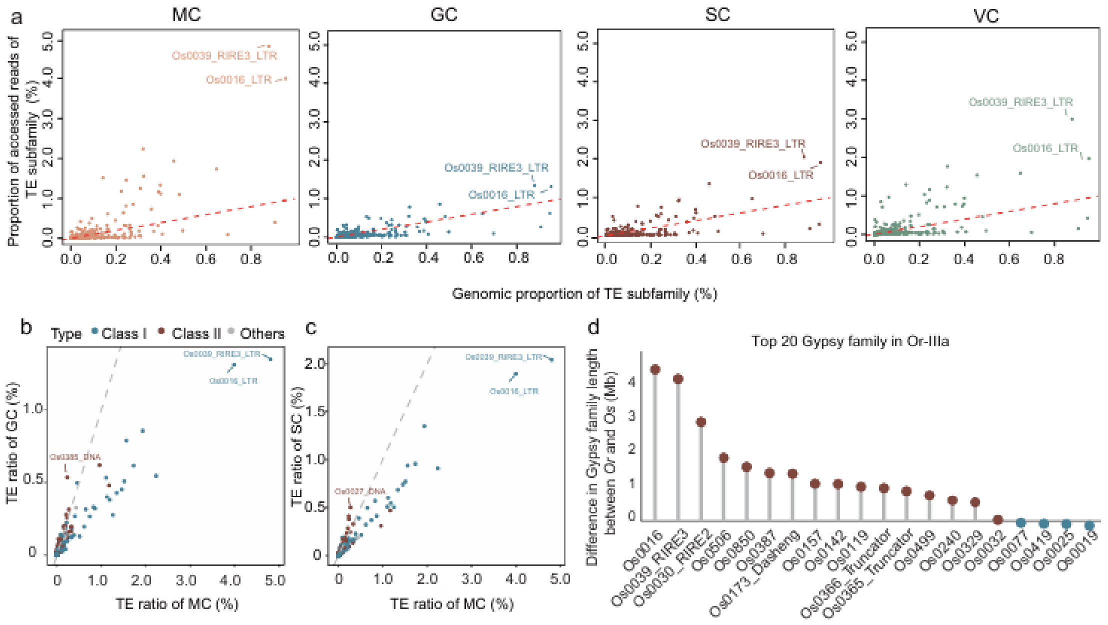


**Fig S3**. **Chromatin accessibility and TE characteristics across cell types**. **a**, Scatter plots showing the correlation between the genomic abundance of each TE subfamily and its chromatin accessibility (proportion of ATAC-seq reads) in MC, GC, SC, and VC. Each dot represents a TE subfamily. Representative subfamilies are labeled. The dashed line indicates equal proportions. **b-c**, Pairwise correlations of accessible TE read ratios between MC and GC (**b**) and between MC and SC (**c**). **d**, Bar plot showing the top 20 *Gypsy* family elements in *O. rufipogon* (Or-IIIa) and the comparison of their total lengths between *O. rufipogon* (Or) and *O. sativa* (Os). Comparative length data were obtained from Guo *et al*., 2025, Extended Data Fig. 2.

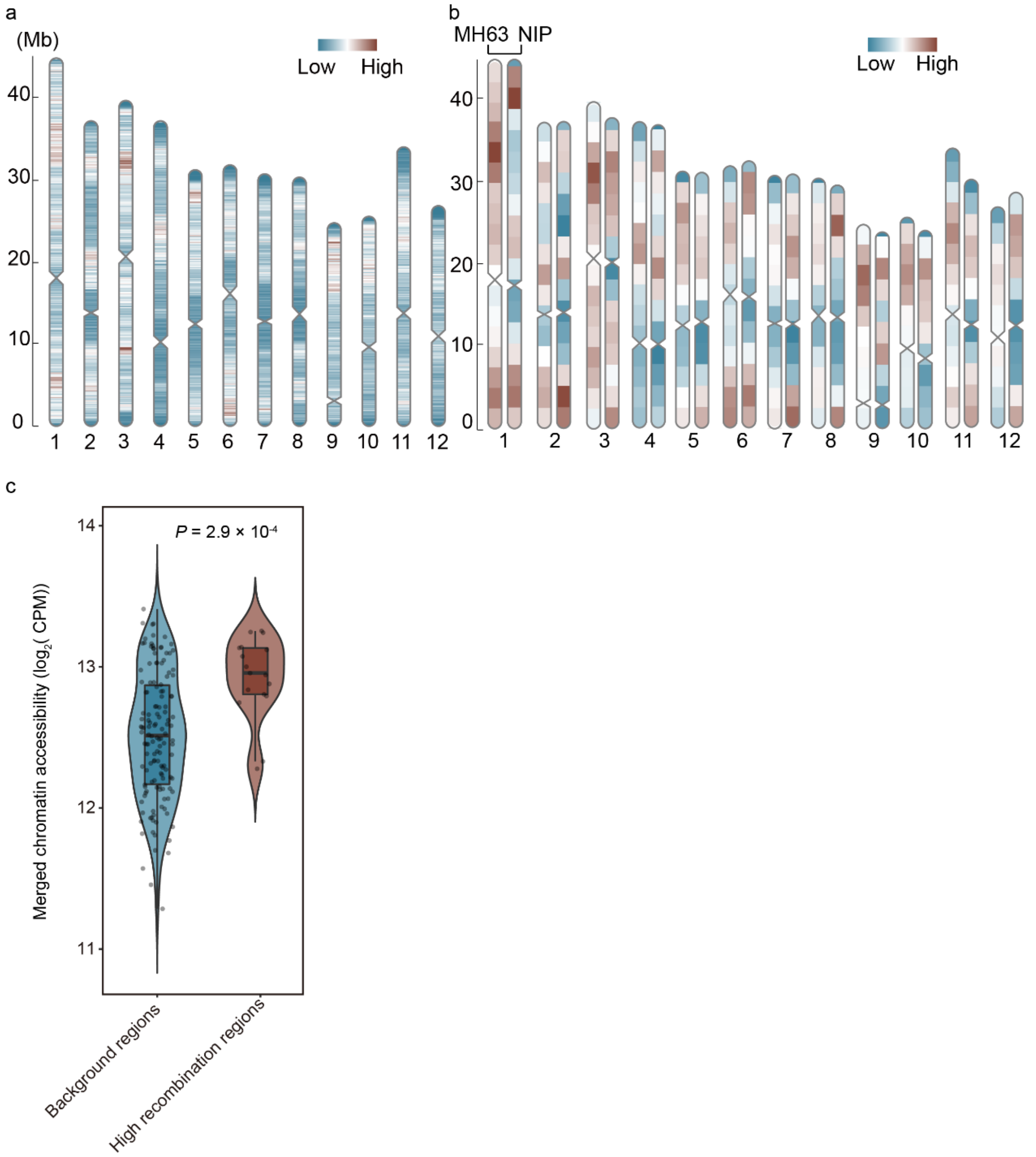


**Fig S4. Genome-wide landscape of recombination events. a**, Distribution of recombination events across the MH63RS3 genome detected in filtered single cells. Events were calculated using 10 kb windows. **b**, Comparison of recombination rate distributions between the single-cell pollen population (this study; mapped to MH63RS3) and a previously reported MH63 × ZS97 recombinant inbred line (RIL) population (Yu *et al.*, 2011; mapped to NIPv6). Recombination rates were calculated using 5 Mb windows with a 2.5 Mb step size. **c**, Violin plots comparing CA between high-recombination and background genomic windows. The genome was partitioned into 2.5-Mb windows; the top 10% of windows by recombination rate were defined as high-recombination regions and the remainder as background. CA was quantified as $\log_2$ (CPM) from merged MPS-ATAC-seq reads across all four cell types; each point is one window. Center lines, medians; boxes, interquartile ranges. Significance was assessed by a two-sided *Wilcoxon* rank-sum test ($P = 2.9 \times 10^{-4}$).

a

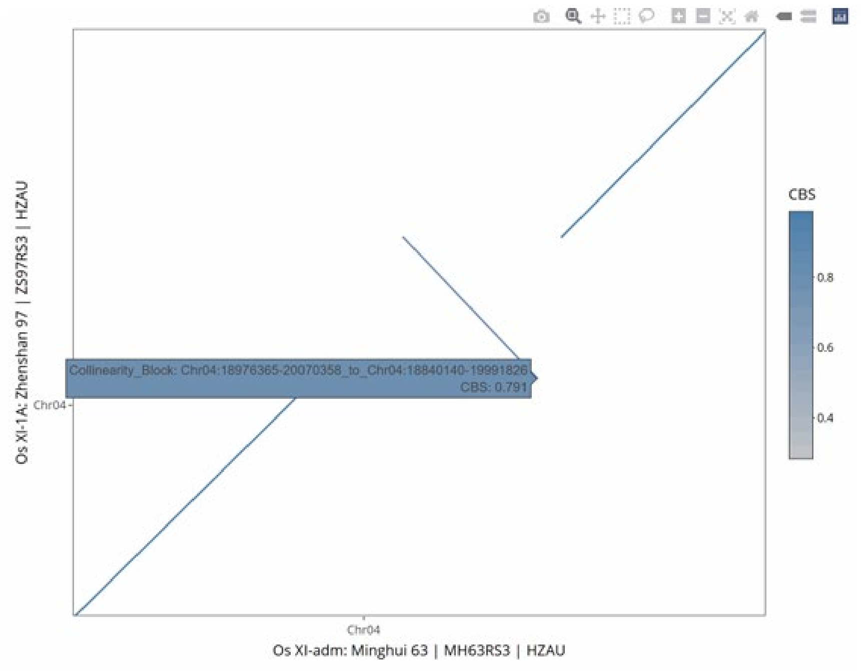


**Fig S5. Synteny analysis between MH63 and ZS97 of the SD loci on chromosome 4**. The dot plot was generated using the MacroCollinearity function of the Rice Gene Index (RGI) (https://riceome.hzau.edu.cn/) illustrates the collinearity between MH63 (x-axis) and ZS97 (y-axis). The diagonal break and reverse slope indicate a large inversion event spanning approximately 18.96-20.06 Mb on chromosome 4. This structural variation is located within the region of observed segregation distortion (SD). Color intensity represents the collinearity clock score (CBS).

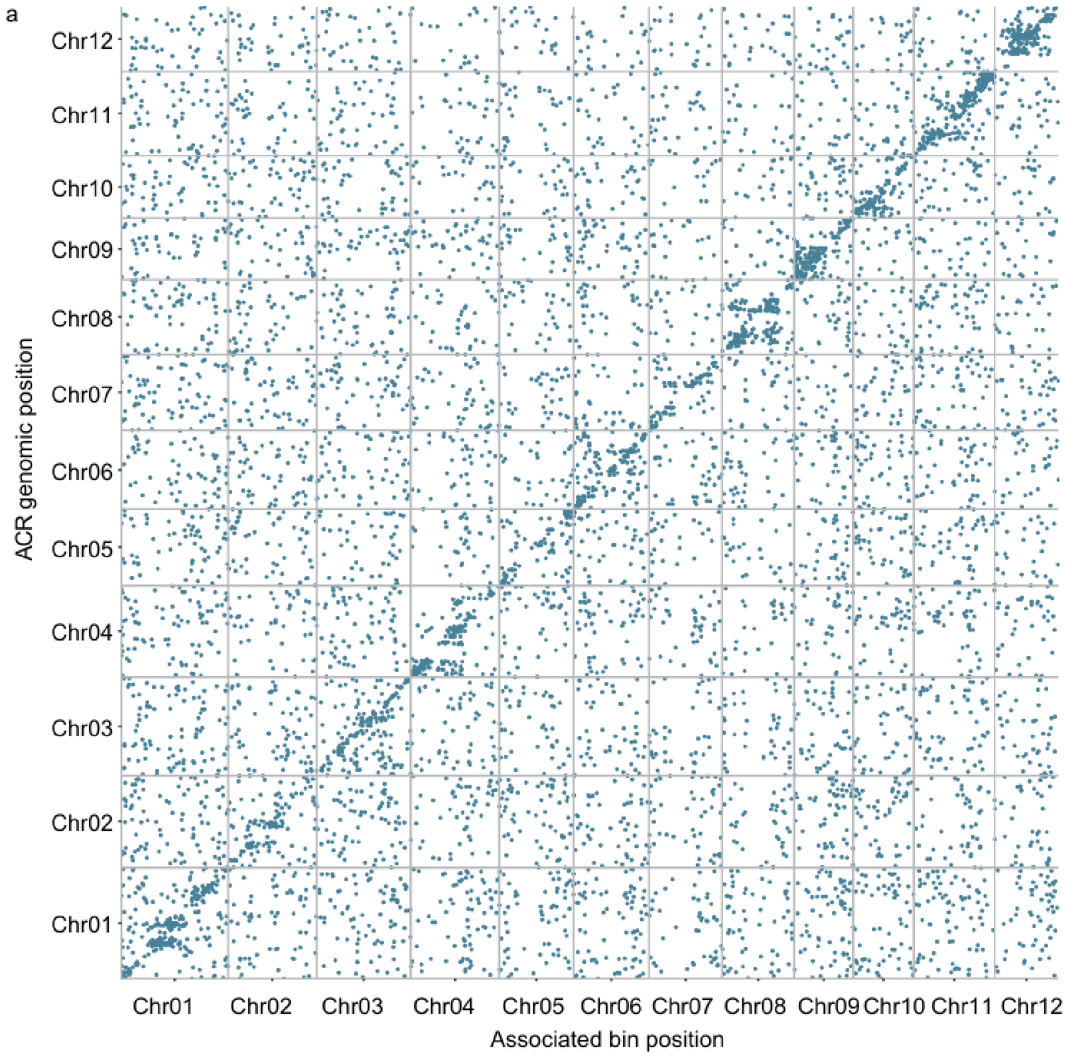


**Fig S6. caQTL distribution at single-cell resolution**. Genome-wide distribution of caQTLs identified by Dynamic-BSA across all nuclei. Each dot represents a significant caQTL association, the x-axis indicates the genomic bin position and the y-axis indicates the ACR position.

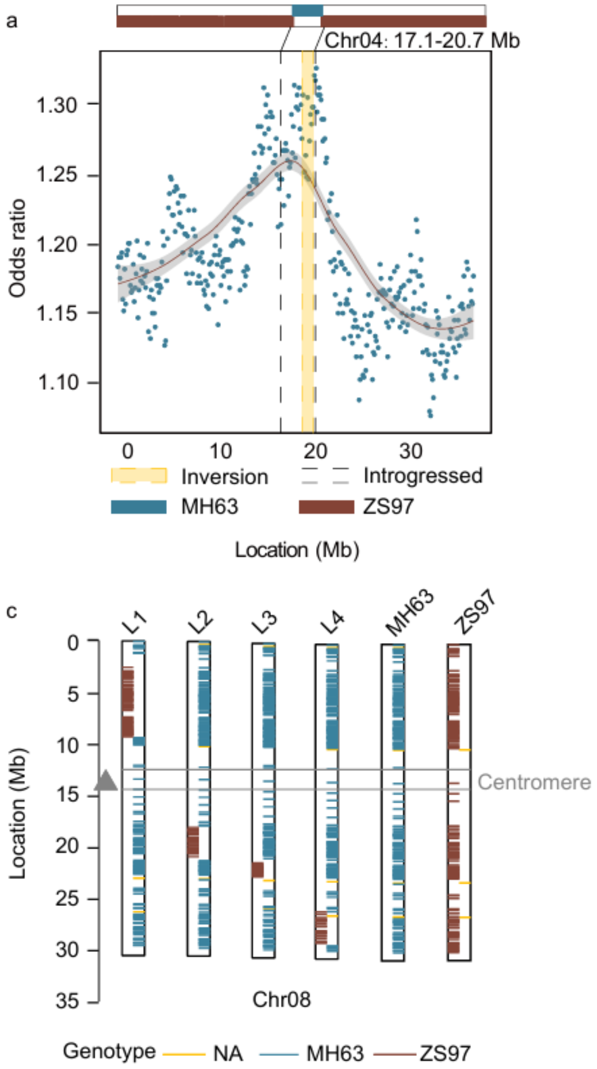


**Fig S7. Validation of Dynamic-BSA and *DTM1* caQTL in VC using introgression lines. a**, Enrichment of DARs within the introgressed regions on chromosome 4. The x-axis represents the positions of the introgressed regions. Blue dots indicating the odds ratio calculated using Fisher's exact test within each 50 kb window, comparing the frequency of DARs between R1 introgression lines and the ZS97 background. The red curve indicates the fitted curve from loess regression. Vertical dashed lines demarcate the boundaries of the introgressed segment (17.1-20.7 Mb) derived from MH63 in the ZS97 background. The red curve shows the fitted trend obtained by loess regression. **b**, Comparison of *P* values for caQTLs identified by the Dynamic-BSA method (genotype-based pool) and those derived from random sampling (random pool). **c**, Genotype verification of the introgression lines used for *DTM1* validation. Each bar represents a genomic variant. Blue bars represent the MH63 genotype, brown bars represent the ZS97 genotype, and yellow bars indicate heterozygous regions or ambiguous genotype calls. The grey triangle and horizontal box mark the centromere position. L1-L4 contain specific ZS97 segments introgressed into the MH63 background: L1 (2.28-9.66 Mb), L2 (17.30-20.87 Mb), L3 (20.87-25.13 Mb), and L4 (25.68-28.80 Mb). MH63 and ZS97 serve as controls (detailed information is provided in Table S4).

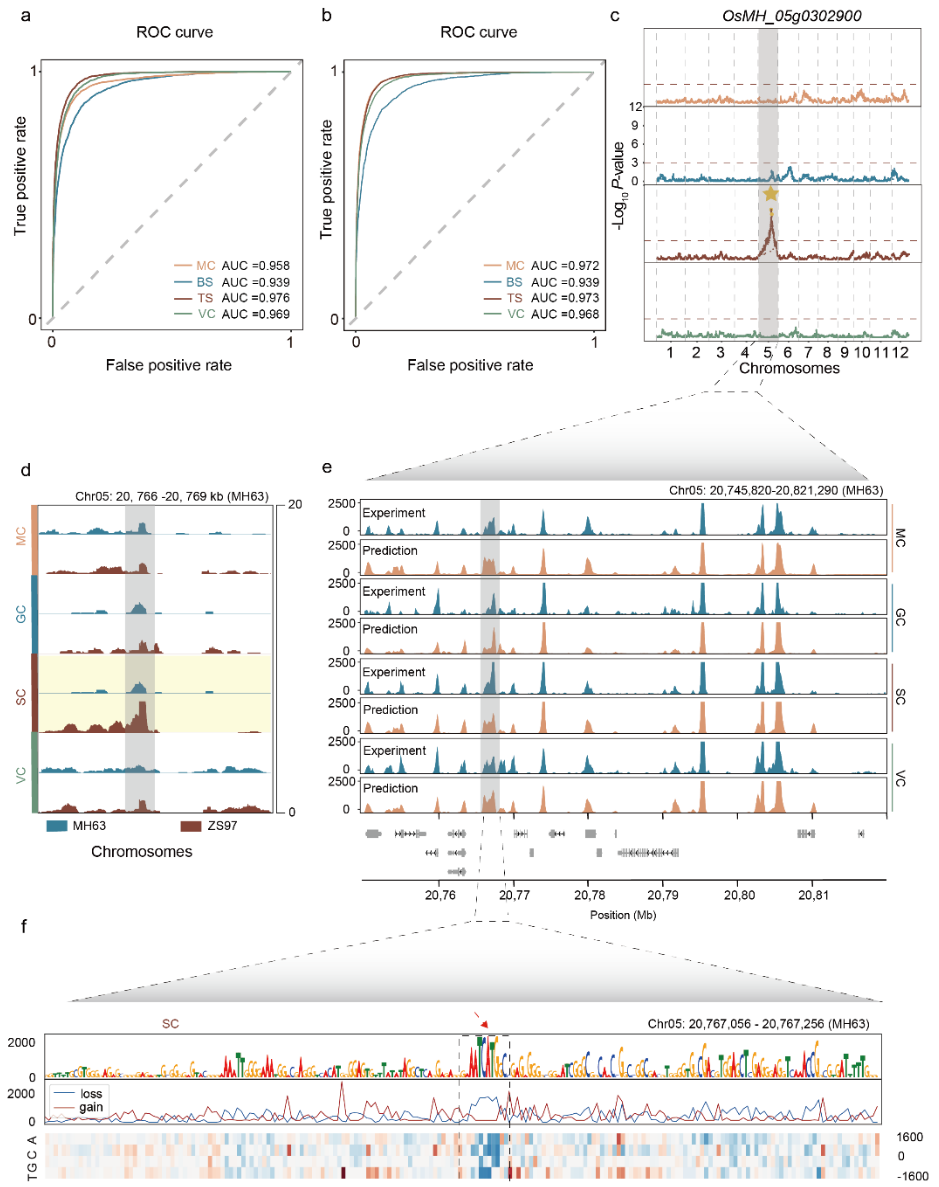


**Fig S8. Interpreting of caQTL variants using a deep learning model**. **a-b**, Performance evaluation of the Basenji model. Receiver operating characteristic (ROC) curves and area under the curve (AUC) scores are shown for models trained on multiple cell types using the MH63RS3 (**a**) and ZS97RS3 (**b**) reference genomes. **c**, Genome-wide caQTL mapping results for the *OsMH_05g0302900* locus across four cell types. The yellow star marks the significant ACR. **d**, Chromatin accessibility profiles of *OsMH_05g0302900* across cell types, with reads separated by genotype (MH63 in blue and ZS97 in brown). **e**, Comparison of experimental (top) and predicted (bottom) chromatin accessibility profiles for the *OsMH_05g0302900* promoter region. **f**, In silico saturation mutagenesis of a 200-bp sequence using the SC model. The heatmap displays predicted accessibility changes for every possible mutation: rows represent the substituted nucleotide, and columns represent genomic positions. Red indicates increased accessibility (gain score), while blue indicates decreased accessibility (loss score). The top track shows the sequence logo derived from loss score, the dashed box highlights a predicted ARR motif disrupted by the variant (indicated by the red arrow).

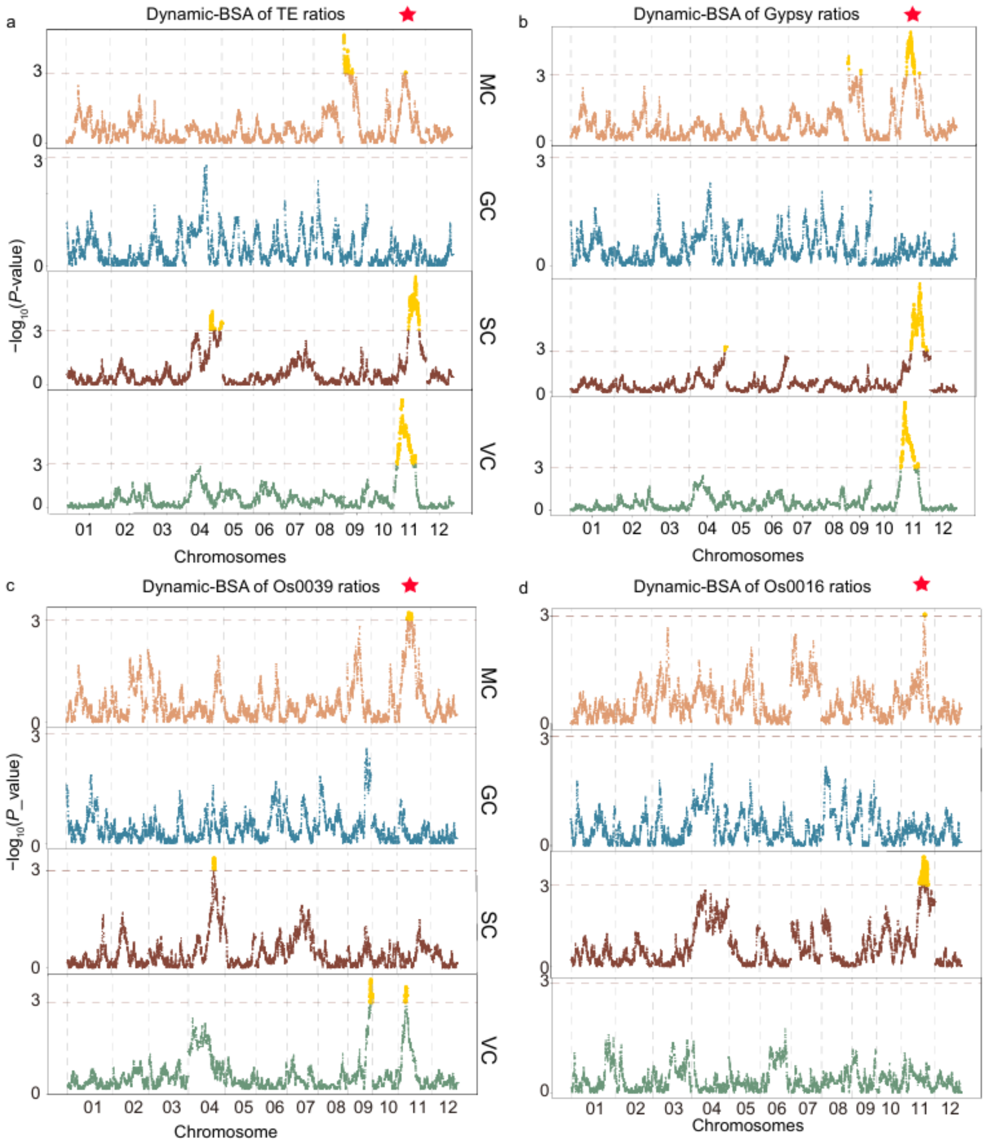


**Fig S9. Genome-wide Dynamic-BSA of TEs and gypsy element ratios**. Each panel displays results for four cell types (from top to bottom: MC, GC, SC, and VC). Red stars indicate the most significant QTL ACR. **a**, Genome-wide Dynamic-BSA of TE ratios. The y-axis indicates the -log$_{10}$($P$ value) for association at each 50 kb window. **b**, Genome-wide Dynamic-BSA of gypsy element ratios. **c,** Genome-wide Dynamic-BSA of Os0039 ratios. **d,** Genome-wide Dynamic-BSA of Os0016 ratios.
